# Timing of DNA damage checkpoint adaptation predicts post-mitotic proliferative competence

**DOI:** 10.64898/2026.09.15.751746

**Authors:** Ambra Dondi, Erika Calabrese, Clara Visintin, Sara Spreafico, Rosella Visintin

## Abstract

Persistent DNA damage creates a conflict between checkpoint-enforced arrest and continued proliferation. Although adaptation enables cells to escape this arrest, checkpoint bypass does not ensure renewed proliferation. Here, we show that the duration of checkpoint-associated metaphase arrest predicts proliferative re-entry after telomere dysfunction in budding yeast. Using population-level analyses and single-cell live imaging, we resolved adaptation into checkpoint attenuation, anaphase entry, mitotic completion and renewed proliferation. These transitions were separable: cells could enter anaphase yet fail to resume proliferation. Among cells that bypassed the checkpoint, each additional 100 min of metaphase arrest was associated with an 80% reduction in the odds of complete proliferative re-entry. This inverse relationship was reproducible across independent experiments and persisted across changes in spindle assembly checkpoint activity, the severity of telomere dysfunction and DNA-end metabolism, despite context-dependent shifts in arrest timing and overall proliferative competence. Comparisons across damage contexts and checkpoint states further showed that the predictive value of metaphase duration depends on the trajectory in which arrest develops. Thus, checkpoint bypass and productive proliferation are separable outcomes, with arrest duration providing a context-dependent predictor of proliferative competence after adaptation.

## Introduction

DNA damage threatens genome integrity by forcing cells to balance repair with the need to preserve proliferative capacity. DNA damage checkpoints delay cell-cycle progression through conserved signaling pathways centered on ATM/ATR-family kinases and downstream checkpoint effectors, thereby coordinating DNA repair with cell-cycle transitions and limiting the propagation of damaged genomes [1–3]. When lesions are successfully repaired, checkpoint signaling is attenuated and cells resume proliferation through checkpoint recovery [4–6]. When damage persists, however, cells can escape prolonged checkpoint arrest through checkpoint adaptation, progressing through mitosis despite unresolved DNA lesions [5–9]. Although adaptation can promote short-term survival, it also allows damaged chromosomes to be transmitted and can thereby contribute to genome instability [6, 7, 10, 11]. This is particularly relevant in cancer, where DNA-damaging chemotherapy and radiotherapy impose chronic genotoxic stress and checkpoint escape or adaptation-like responses may allow tumor cells to retain proliferative capacity, contributing to treatment resistance at the cost of increased genome instability [11–13].

Checkpoint adaptation has been described from budding yeast to mammalian cells and is increasingly viewed as a regulated response to chronic genotoxic stress rather than a passive collapse of checkpoint control [6, 8–10, 14, 15]. The logic by which adaptation produces different cellular outcomes remains incompletely understood. Adaptation has traditionally been inferred from endpoint measurements such as checkpoint kinase phosphorylation, rebudding, colony formation, or cell morphology [16]. These assays are informative at the population level, but they do not distinguish checkpoint attenuation from anaphase entry, successful mitotic completion or subsequent proliferative re-entry. As a result, mechanistically distinct states can appear phenotypically equivalent, and adaptation is often treated as a binary arrest-versus-escape decision. Single-cell studies in mammalian systems have revealed that DNA damage responses are highly dynamic and that temporal features of checkpoint signaling can influence cell fate [17–19]. In human cells, outcomes following DNA damage-induced arrest reflect the balance between residual checkpoint signaling and pro-mitotic activities, while checkpoint dynamics after replication stress influence daughter-cell fate, including p53/p21-dependent arrest after mitotic passage [17, 20–22].

Mitotic duration itself is increasingly recognized as a biologically encoded signal rather than a passive consequence of cell-cycle perturbation. In mammalian cells, surveillance pathways can convert prolonged mitotic residence into p53-dependent post-mitotic arrest, thereby limiting the propagation of cells that divide outside the normal temporal range [23–26]. These findings have led to the concept of mitotic “clocks” that couple elapsed mitotic time to subsequent cell fate and genome protection [27]. Human cells can also escape a sustained G2 checkpoint and enter mitosis with persistent DNA damage, producing outcomes that include mitotic death and survival with micronuclei [14, 28]. Yet whether time spent in the preceding damage-induced arrest predicts which adapted cells retain proliferative competence remains unknown. This raises the possibility that adaptation outcome depends not only on whether cells bypass the checkpoint, but also on when they do so and whether they retain the capacity to execute mitosis and resume proliferation. Persistent damage may therefore generate intermediate states in which checkpoint activity, mitotic competence and proliferative potential become progressively uncoupled.

Budding yeast provides a powerful system for addressing this question because checkpoint adaptation can be monitored in individual cells following sustained activation of the DNA damage checkpoint. Two complementary approaches are commonly used to trigger this response. The temperature-sensitive *cdc13-1* allele induces telomere uncapping at the restrictive temperature and has been widely employed to study adaptation to persistent chromosome-end dysfunction [29–31]. In contrast, inducible *HO* endonuclease cleavage activates the checkpoint through the generation of a defined DNA double-strand break [32]. These systems can also be combined with genetic perturbations affecting checkpoint persistence, spindle assembly checkpoint activity and DNA-end metabolism, allowing these variables to be analyzed without assuming that endpoint adaptation phenotypes reflect a single underlying state.

Here, we integrate population-level analyses with single-cell live imaging to distinguish checkpoint attenuation, anaphase entry, mitotic execution and renewed proliferation. Using persistent telomere dysfunction as a heterogeneous damage context, we ask whether cell-to-cell variation in metaphase arrest duration merely reflects asynchronous checkpoint escape or instead predicts subsequent fate. We show that checkpoint bypass is necessary but not sufficient for productive proliferation: metaphase duration quantitatively orders post-adaptation outcomes across independent *cdc13-1* datasets and genetic backgrounds, with prolonged arrest associated with a progressive decline in complete proliferative re-entry. Perturbations affecting spindle assembly checkpoint activity, the severity of telomere dysfunction and DNA-end metabolism shift arrest timing without eliminating this inverse timing-outcome relationship. Increased damage severity and altered DNA-end metabolism also change, in a context-dependent manner, the baseline probability of productive re-entry, whereas cells responding to a single *HO*-induced double-strand break occupy a narrower temporal range in which robust proliferation predominates. Checkpoint-null *rad9Δ* and delayed-checkpoint *exo1Δ* states further show that checkpoint state determines how metaphase duration should be interpreted. Together, these findings define checkpoint adaptation as a temporally structured cell-fate trajectory in which the consequences of checkpoint escape depend not only on whether damaged cells divide, but also on when they do so and whether they remain competent to complete mitosis and resume proliferation.

## Materials and Methods

### Yeast strains

All *cdc13-1* mutant strains were derived from the W303 (K699) background and are *RAD5^+^*, whereas the *GAL-HO* strains were derived from JKM179. The relevant genotypes of the strains used in this study are listed in Supplementary Table 1.

### Growth conditions

Cell cycle synchronization experiments were performed as described previously [33]. *cdc13-1* strains were grown at 23°C in yeast extract peptone medium supplemented with 2% glucose (YEPD); whereas *GAL-HO* strains were grown in yeast extract-peptone medium supplemented with 2% raffinose (YEPR). Cells were arrested in G1 phase using 5 μg/ml α-factor (GenScript). For population analyses, G1-arrested cultures were washed with 10 volumes of fresh medium lacking pheromone. For single-cell imaging, cells were washed and loaded directly into Y04C CellASIC microfluidic plates containing the appropriate release medium. *cdc13-1* cells were released into YEPD at 32°C, whereas *GAL-HO* cells were released at 23°C into YEPR supplemented with 2% galactose to induce HO expression.

### Immunoblot analysis

Cells were treated with cold 5% TCA for 10 min, pelleted, and washed with acetone. Pellets were resuspended in 50 mM Tris-HCl (pH 7.5), 1 mM EDTA, 1 mM p-nitrophenyl phosphate, 50 mM DTT, 1 mM PMSF, and 2 μg/ml pepstatin, lysed with glass beads, and boiled in 1x SDS sample buffer. Proteins were detected with the appropriate antibodies.

### Antibody information

The primary antibodies used were EL7.E1 monoclonal anti-Rad53 (gift from Dr. M. Foiani [34]) to detect total Rad53 protein, F9 monoclonal anti-phosphorylated Rad53 (gift from Dr. M. Foiani [34]), anti-Kar2 (gift from Dr. J. V. Kilmartin) to detect Kar2 protein. The secondary antibodies used were Goat anti-rabbit IgG (H + L)-HRP conjugate (170-6515, Bio-Rad, used at 1:10,000 dilution) and goat anti-mouse IgG (H + L)-HRP conjugate (170-6516, Bio-Rad, used at 1:10,000 dilution). Proteins were visualized by chemiluminescence (ECL, GE Healthcare).

### DNA content analyses using flow cytometry

Cell culture samples (1 ml, OD600 = 0.2 - 0.4) were collected by centrifugation for 1 minute at 13 000 rpm at RT and incubated O/N at 4°C in 70% ethanol. The cells were then washed with 1 ml 50 mM Tris-HCl pH 7.4 and incubated O/N at 37°C in the same buffer containing 1 mg/ml RNase A. Next, the cells were collected by centrifugation, resuspended in 1 ml 55mM HCl containing 5mg/ml pepsin (Sigma) and incubated at 37°C. After 30 minutes, the cells were collected, resuspended in 50 mM Tris-HCl pH 7.4 and sonicated 3 times for 10 seconds with intervals of 30 seconds using a Bioruptor UCD-300 (Diagenode) water-bath sonicator set on low intensity. Immediately before FACS reading, 100 ul of cell suspension were added to 1 ml of Sytox Green staining solution and the samples were acquired with FACSCalibur system (Becton Dickinson) operated via the CellQuest software. The data were analyzed with FlowJo Analysis 8.8.6 software.

### Microscopy, live-cell imaging and image analysis

*For population experiments*: cells were fixed with 4% paraformaldehyde to visualize spindle morphology and Scc1 dynamics. Nuclear morphology was assessed by DAPI staining using mounting solution containing 0.04 M K₂HPO₄, 0.01 M KH₂PO₄, 0.15 M NaCl, 0.1% NaN₃, 0.05 μg/ml DAPI, 0.1% p-phenylenediamine and 90% glycerol. Fixed cells were imaged with GFP, mCherry, and DAPI filter sets, and bright-field mode using an Andor Zyla VSC-04470 sCMOS camera mounted on a Leica DM6 B MultiFluo microscope equipped with a Plan-Apochromat 100X/1.4 NA objective. No binning, z=10 stacks (0.5μm step size).

*n* = 100 cells were analyzed *per* timepoint *per* strain. Cells were classified as interphase when they contained interphase microtubules, including G1, S and G2 cells; metaphase when they contained short bipolar spindles and/or persistent Scc1 signal; and anaphase when they contained elongated mitotic spindles accompanied by loss or redistribution of Scc1 signal. Budding index was scored in parallel by cell morphology. Cells were classified as unbudded, large-budded dumbbell cells, or rebudded cells. Rebudded cells were defined as previously arrested dumbbell cells in which one or both compartments had produced a new bud. Images were analyzed using Fiji.

*For single-cell experiments:* Live-cell fluorescence imaging was performed using either a DeltaVision Elite deconvolution microscope (Applied Precision) equipped with an Olympus IX71 inverted microscope, an Olympus UPLS Apo 100X/1.40NA, and a CoolSNAP HQ2 CCD camera (Photometrics), or a THUNDER Imager Cell widefield fluorescence microscope (Leica Microsystems) based on a fully motorized DMi8 inverted microscope, equipped with a Leica HC PL APO 100X/1.40NA objective and a Leica DFC9000 GTC camera. DeltaVision image acquisition was controlled using SoftWoRx software, whereas THUNDER imaging was controlled using LAS X software.

For live-cell imaging, GFP and mCherry fluorescence signals were acquired using the corresponding fluorescence filter sets together with DIC reference images. Five z-sections spaced 1 µm apart were acquired every 15 min for up to 20 h. DeltaVision images were acquired using 2 x 2 binning, and focus was maintained throughout the time lapse using the Ultimate Focus hardware autofocus system. Images were subsequently deconvolved using SoftWoRx software. For THUNDER acquisitions, images were collected using 2 x 2 binning, and focus was maintained throughout the time lapse using Leica Adaptive Focus Control (AFC). THUNDER fluorescence images were processed using THUNDER Computational Clearing to reduce out-of-focus fluorescence.

Time-lapse image sequences were analyzed using Fiji/ImageJ. Mitotic progression was followed using fluorescent markers of spindle dynamics and cell-cycle progression, and DIC images were used to monitor cell morphology and budding. Metaphase duration was determined from bipolar spindle formation to spindle elongation/disassembly and Scc1 disappearance. Post-mitotic proliferative re-entry was assessed by monitoring rebudding of the resulting cellular compartments.

*n* = cell number specified directly in Figures.

*For budding only experiments* (pertinent to Supplementary Figure 1 only): Images were acquired with an Olympus U Plan Apo 40X dry objective (NA 0.85) every 15 minutes for 20 hours.

### Statistics and data analysis

Unless otherwise indicated, statistical analyses were performed using single-cell measurements. To quantify metaphase arrest, we defined metaphase duration as the interval between bipolar spindle formation and anaphase onset, identified by spindle elongation and/or disassembly together with cohesin cleavage, marked by Scc1 disappearance. Because Scc1 loss consistently coincided with the corresponding change in spindle dynamics, in a subset of experiments spindle behavior was used alone. Spindle dynamics and Scc1 signal were measured from time-lapse images acquired at 15-min intervals. Because a metaphase-exit time could be assigned only to cells that underwent Scc1 loss and/or elongated their spindles, analyses of metaphase-duration distributions and duration-dependent outcomes were conditional on anaphase entry. Cells that remained arrested or died before anaphase entry were analyzed separately as non-adapting cells and were not assigned a metaphase duration for duration-based analyses or predictive modeling. No assumption of normality was made.

Post-mitotic rebudding was scored for the two compartments of the arrested mother-daughter pair as follows: score 0, neither compartment rebudded; score 1, one compartment rebudded; and score 2, both compartments rebudded. All cells with an observed metaphase-exit time and a scorable post-mitotic outcome were included in analyses relating metaphase duration to rebudding score. No cells were excluded from these analyses on the basis of extreme metaphase-duration values.

Associations between metaphase duration and rebudding outcome were assessed using Spearman’s rank correlation coefficient because rebudding score is ordinal and metaphase duration was not assumed to follow a normal distribution. Spearman’s ρ and the corresponding two-sided *P* values were calculated for individual datasets and for pooled analyses. For pooled analyses, single-cell measurements from independent experiments were combined. All cells were retained in the correlation analyses, including those with extreme metaphase-duration values.

Metaphase-duration distributions were compared across experiments or genotypes using the Kruskal-Wallis test. When the omnibus test was significant, pairwise comparisons were performed using two-sided Mann-Whitney U tests with Holm correction for multiple testing. For comparisons involving two groups, a two-sided Mann-Whitney U test was used directly. These analyses correspond to the metaphase-duration distributions presented as violin plots and are reported in the relevant figure legends and Results sections. Violin plots show kernel-density estimates, with individual cells overlaid as points. Central tendency and dispersion are summarized by the median and interquartile range.

Sensitivity analyses excluding extreme metaphase-duration values were performed only for the specified comparisons of metaphase-duration distributions. Outliers were identified using Tukey’s 1.5 x interquartile-range criterion, applied independently within each individual dataset for dataset-level analyses and within each pooled genotype for genotype-level comparisons. Analyses relating metaphase duration to rebudding outcome used the complete datasets. As a sensitivity analysis, the primary binary logistic model was also repeated after within-dataset exclusion of values identified using Tukey’s 1.5 x IQR criterion (Supplementary Table 2); all other duration–outcome analyses retained the complete datasets. For selected pooled scatter-plot visualizations, cells with metaphase durations greater than 800 min were omitted solely to prevent compression of the main dynamic range. These cells were retained in all corresponding statistical analyses unless explicitly stated otherwise.

Statistical significance was defined as *P* < 0.05. Analyses and data visualization were performed in Python using SciPy and Matplotlib.

### Predictive modeling of post-adaptation proliferative outcome

Single-cell measurements were analyzed in R using logistic regression models to test whether metaphase arrest duration predicted post-adaptation proliferative outcome. Metaphase duration was defined as the interval from bipolar spindle formation to anaphase onset, identified by Scc1 loss together with spindle elongation, or from spindle dynamics alone in datasets in which Scc1 was not imaged. Proliferative outcome was scored after mitotic exit according to rebudding of the two compartments of the arrested mother-daughter pair: 0, neither compartment rebudded; 1, one compartment rebudded; 2, both compartments rebudded. Full model specifications, model evaluation and transfer procedures and refitting details are provided in the Supplementary Method. For the primary analysis, proliferative outcome was converted to a binary variable. Cells with a score of 2 were classified as full proliferative capacity, whereas cells with scores of 0 or 1 were classified as incomplete or failed proliferative competence. Metaphase duration was included as a continuous predictor and scaled to 100-min intervals so that model coefficients represent the effect of each additional 100 min spent in metaphase arrest. Where multiple experiments were analyzed together, dataset identity was included as a covariate to account for experiment-to-experiment variability in absolute timing or productive adaptation frequency.

For genotype comparisons, two complementary logistic-regression models were fitted. An additive model containing metaphase duration and genotype, together with dataset or experimental-block terms where appropriate, tested the duration-adjusted genotype effect. A second model included a metaphase duration x genotype interaction to test whether the duration-outcome slope differed between genotypes. A nonsignificant interaction indicated no evidence of a difference between genotype-specific slopes and was not interpreted as evidence of equivalence. Model outputs, including coefficients, confidence intervals, p-values, and performance metrics, are reported in Supplementary Table 2. Sample sizes and the role of each dataset group in these analyses are listed in Supplementary Table 3.

As a complementary analysis, the full 0-1-2 proliferative outcome score was analyzed using ordinal logistic regression. Temporal thresholds were used only for visualization and descriptive stratification and were not used for primary statistical inference.

## Results

### Single-cell analysis separates checkpoint bypass from productive proliferation

Conventional assays of checkpoint adaptation rely largely on endpoint readouts such as Rad53 phosphorylation, morphology, rebudding or colony formation [4, 8, 9, 16, 35]. To reconstruct adaptation as a dynamic process, we combined population-based analyses with single-cell live imaging in a multiparameter framework that distinguishes checkpoint signaling, mitotic progression and proliferative re-entry across complementary population-level and single-cell readouts (Fig. 1A). This approach allowed us to separate four transitions that are often conflated: checkpoint attenuation, measured at the population level by declining Rad53 phosphorylation; checkpoint bypass, defined as anaphase entry despite persistent DNA damage; mitotic execution, assessed by anaphase progression and completion of division; and post-adaptation proliferative re-entry, defined by rebudding.

**Figure 1.**
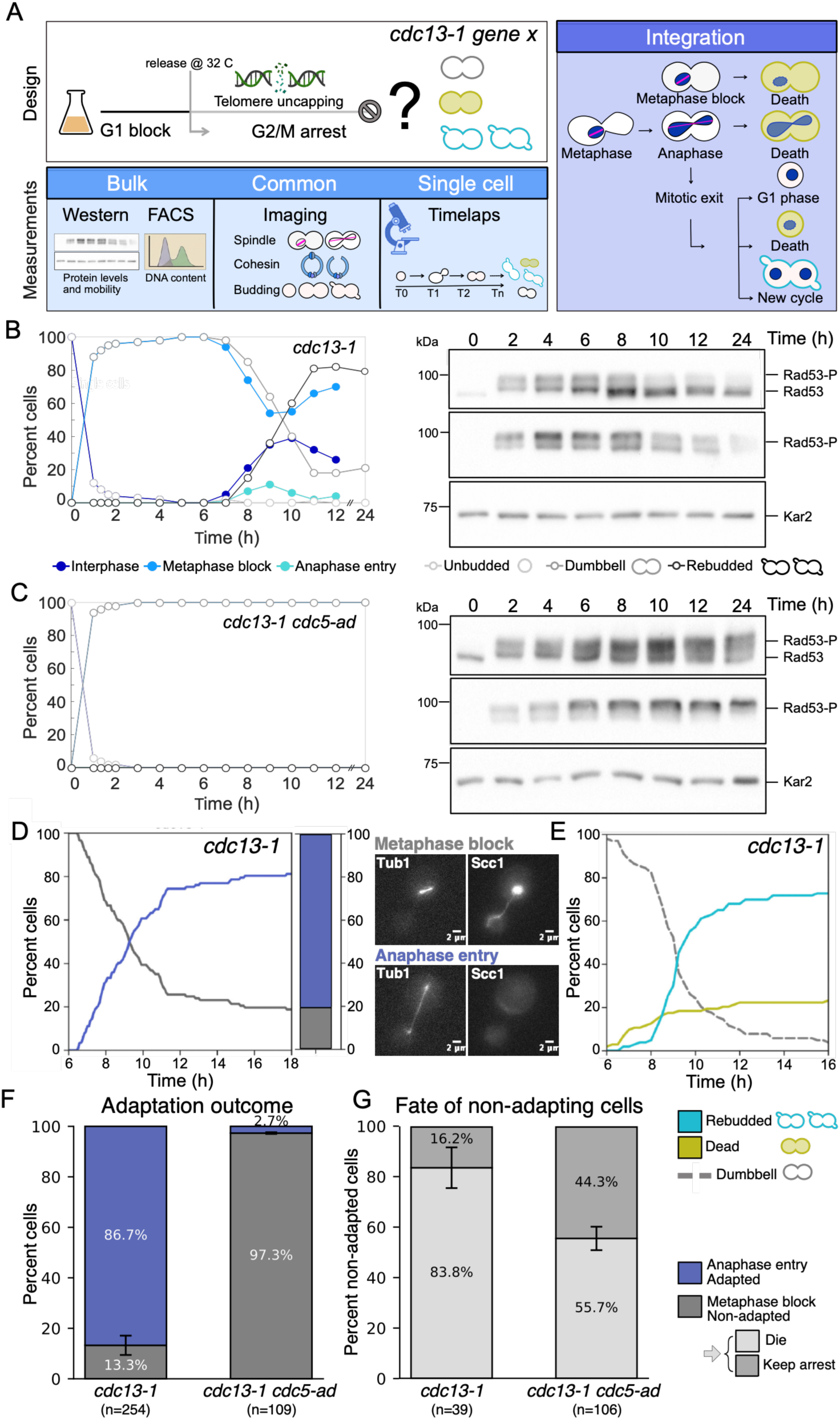
Single-cell tracking links DNA damage checkpoint adaptation to cell-fate outcomes after telomere uncapping. **(A)** Experimental workflow for analyzing mitotic fate after telomere uncapping in *cdc13-1* cells. Cells were released from G1 arrest at 32°C to induce DNA damage and followed by bulk assays, microscopy, and single-cell time-lapse imaging. **(B-C)** Bulk time-course analysis of *cdc13-1* (Ry9666, **B**) and *cdc13-1 cdc5-ad* (Ry10642, **C**) cells carrying *mCherry-TUB1* and *SCC1-yEGFP* fusions after release into telomere-uncapping conditions. Samples were collected at the indicated times and analyzed through fixed cells (Tub1 and Scc1) to monitor mitotic spindle dynamics, Scc1 localization and nuclear morphology. *n*=100 cells *per* timepoint (left graph). Representative immunoblots show Rad53 phosphorylation as a marker of checkpoint activation. Kar2 is used as a loading control (right panel). **(D-E)** Representative single-cell imaging time course of *cdc13-1* (Ry9666) cells carrying *mCherry-TUB1* and *SCC1-yEGFP* fusions. Cells were classified as metaphase-blocked or anaphase-entering after checkpoint activation based on spindle/nuclear morphology and Scc1 localization (**D**). Cells were tracked over time for morphology based on budding index and classified as dumbbell, cell death, or rebudding (**E**). *n* = 100 cells per timepoint. **(F-G)** Quantification of adaptation outcome and fate of non-adapting cells, in *cdc13-1* and *cdc13-1 cdc5-ad* cells following DNA damage. The stacked bars show the percentage of cells entering anaphase or remaining in metaphase block, and the fate of non-adapted/metaphase-arrested cells. Bars represent mean ± SEM from four independent datasets.

Cells carrying the temperature-sensitive *cdc13-1* allele, which impairs telomere capping at restrictive temperature, were synchronized in G1 and released at 32°C, a well-established condition for inducing telomere dysfunction and studying checkpoint adaptation [29–31, 35, 36]. Checkpoint signaling was monitored by Rad53 phosphorylation, whereas cell-cycle progression was assessed by DNA content, spindle morphology, cohesin dynamics, and budding index. In parallel, live-cell imaging enabled continuous tracking of spindle dynamics, cohesin signal, and budding index in individual cells (Fig. 1A). As reference conditions, we compared adapting *cdc13-1* cells, with *cdc13-1 cdc5-ad* cells, which carry an adaptation-defective allele of the polo-like kinase Cdc5 and remain arrested in response to persistent telomere damage [8]. Recent work further showed that recruitment of Cdc5 to spindle pole bodies and phosphorylation of spindle-pole-body components are required for efficient adaptation, identifying these structures as signaling platforms for Cdc5 during persistent DNA damage [37]. Following release at 32°C, both strains activated the DNA damage checkpoint, as shown by Rad53 phosphorylation within 2 h (Fig. 1B-C and Supplementary Fig. 1A-B). Thereafter, however, their trajectories diverged.

In *cdc13-1* cells, Rad53 phosphorylation progressively declined from approximately 6-7 h after release (Fig. 1B and Supplementary Fig. 1A), coinciding with a reduction in metaphase spindles (Supplementary Fig. 1A). Consistent with checkpoint bypass and subsequent cell-cycle re-entry, cells progressively rebudded, reaching approximately 60-80% depending on the experiment (Fig. 1B and Supplementary Fig. 1A). DNA content analysis further confirmed progression beyond G₂/M, with the appearance of cells entering a subsequent G1 phase as well as populations with 3C-4C DNA content, likely reflecting incomplete cytokinesis after adaptation (Supplementary Fig. 1A).

By contrast, *cdc13-1 cdc5-ad* cells maintained high Rad53 phosphorylation and persistent dumbbell morphology (large-budded mother-daughter pairs connected by a narrow neck) and metaphase spindles, with little evidence of productive mitotic progression (Fig. 1C and Supplementary Fig. 1B). Although the frequency of metaphase spindles occasionally declined in this mutant, the absence of subsequent rebudding, together with persistent G₂/M DNA content, suggests that this reflected loss of spindle integrity during prolonged arrest rather than checkpoint bypass (Supplementary Fig. 1B).

### Spindle and cohesin dynamics reveal divergent adaptation fates concealed by morphology

To assess cell fate after checkpoint arrest, we followed individual cells by time-lapse microscopy in an assay functionally analogous to a microcolony analysis, monitoring whether cells could resume successive divisions after DNA damage [4, 8, 9]. In *cdc13-1* cells, approximately 70% of cells successfully rebudded, whereas around 20% died, and 10% remained with a dumbbell morphology. By contrast, a large fraction of *cdc13-1 cdc5-ad* cells failed to resume proliferation, either dying or remaining persistently arrested as dumbbell-shaped cells (Supplementary Fig. 1C). Although these analyses distinguish survival from death, morphology alone did not resolve the underlying adaptation state. Dumbbell-shaped cells can correspond to several distinct cell-cycle states, including checkpoint-arrested metaphase cells, cells that have bypassed the checkpoint and entered anaphase, or post-mitotic cells that have failed to complete cytokinesis (Supplementary Fig. 1D). Importantly, changes in morphology lagged behind biochemical and cytological markers of mitotic progression, reflecting the time required to complete anaphase, cytokinesis and enter a new division cycle (Supplementary Fig. 1A and Fig. 1B).

To define these transitions more precisely, we integrated spindle dynamics with Scc1 signal, used as a marker of cohesin dynamics, in both population-based and single-cell analyses. Persistent Scc1 signal together with a short bipolar spindle identified *bona fide* metaphase block, whereas loss of Scc1 accompanied by spindle elongation or spindle disassembly indicated anaphase entry, hence checkpoint bypass (Fig. 1D) [38, 39]. Accordingly, cells were classified as metaphase-arrested/non-adapted when Scc1 and short bipolar spindles persisted, and as bypassed/adapted when Scc1 loss was accompanied by spindle elongation or spindle disassembly (Fig. 1D). In parallel, cells were scored for viability and budding index. For the budding index, cells were classified as either dead, dumbbell or rebudded. Cells were classified as morphologically dead when they appeared markedly shrunken or collapsed, showed loss of cellular integrity, and lacked detectable fluorescence signal. Rebudded cells were defined as dumbbell cells in which at least one of the two connected cell bodies had produced a new bud, resulting in structures with three or four connected cells (Fig. 1E). Using these criteria, the decline in Rad53 phosphorylation in *cdc13-1* cells correlated with a progressive loss of metaphase-arrested cells and the accumulation of cells entering anaphase, which reached approximately 80-90% of the population (Fig. 1B). In parallel, single-cell analysis scored approximately 80-85% of *cdc13-1* cells entering anaphase (Fig. 1D and F, Supplementary Video 1), with only about 70% completing mitosis and re-entering the cell cycle, indicating that 10-20% of cells failed after checkpoint bypass (Fig. 1E). Consistent with previous studies, checkpoint bypass was nearly abolished in *cdc13-1 cdc5-ad* cells, with only approximately 2.7% of cells entering anaphase (Fig. 1F and Supplementary Video 2) [8, 9, 35–37]. Adaptation frequencies measured by live-cell imaging only marginally differed from population-level measurements, indicating that imaging conditions did not significantly alter adaptation kinetics (Fig. 1B vs 1D-F). Among *cdc13-1* cells that failed to adapt, approximately 80% died during prolonged metaphase arrest, whereas a smaller fraction remained arrested until the end of the 20 h experiment (Fig. 1G). Conversely, in *cdc13-1 cdc5-ad* cells, non-adapting cells more frequently remained arrested and showed comparatively less cell death (Fig. 1G). Together, these findings indicate that anaphase entry marks escape from checkpoint arrest but is not sufficient to ensure productive proliferation. Checkpoint bypass and proliferative competence are therefore separable outcomes, and cells that bypass the checkpoint can follow distinct post-arrest trajectories that remain obscured when adaptation is inferred from morphology alone.

### Metaphase arrest duration predicts proliferative competence after checkpoint bypass

This separation between checkpoint bypass and proliferative competence prompted us to ask whether the duration of the preceding metaphase arrest distinguished bypass events that led to productive proliferation from those that did not. To quantify metaphase arrest, we defined metaphase duration as the interval between bipolar spindle formation and anaphase onset, identified by spindle elongation together with cohesin cleavage, marked by Scc1 disappearance. Among cells that entered anaphase, metaphase duration varied widely, ranging from less than 2 h to more than 15 h, with most cells arresting for approximately 4-10 h and an average duration of around 7-8 h (Fig. 2A). Thus, successful checkpoint bypass in *cdc13-1* cells occurs after highly heterogeneous periods of metaphase arrest.

**Figure 2.**
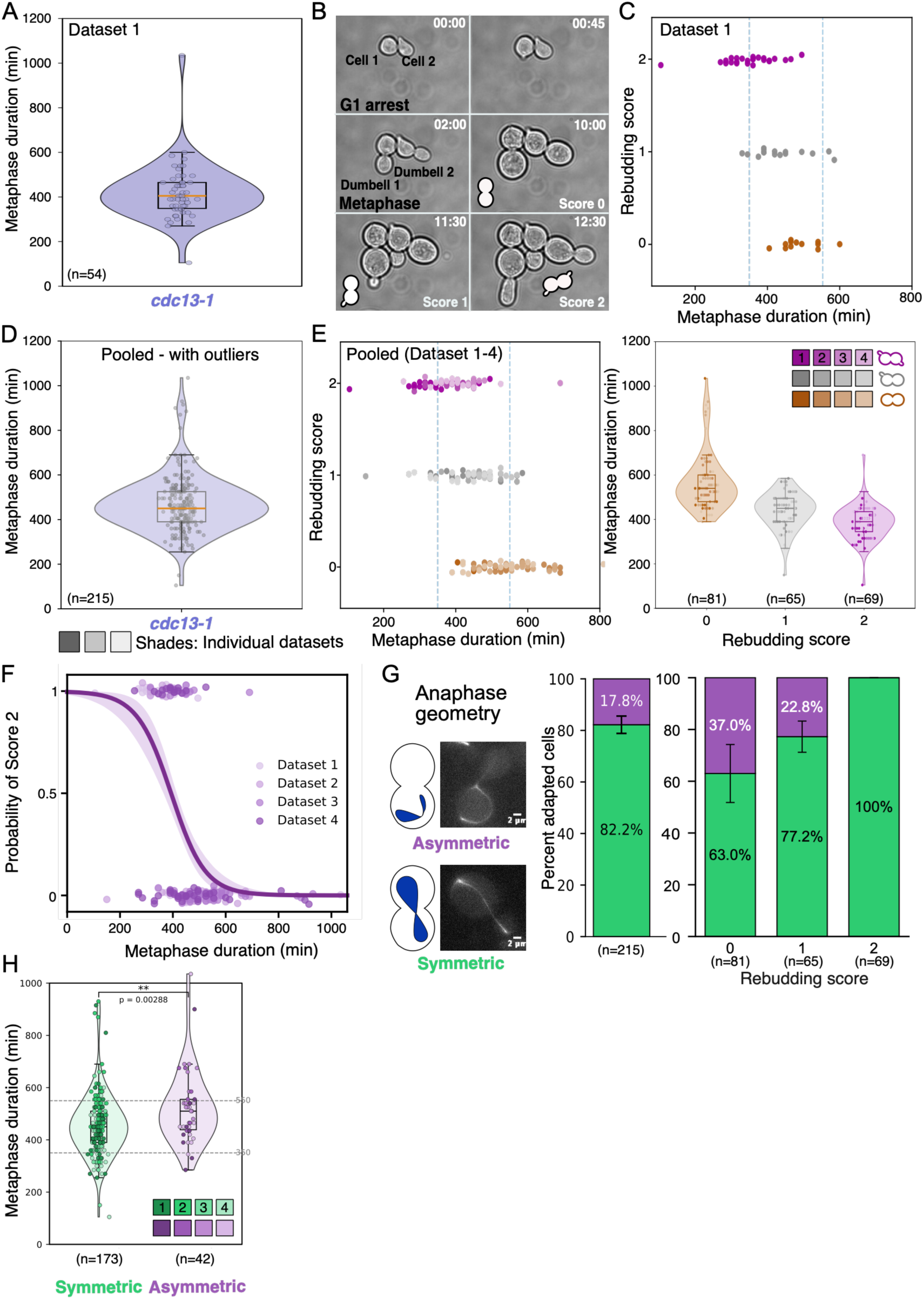
Metaphase duration predicts rebudding outcome and anaphase geometry after telomere uncapping. Synchronously released *cdc13-1* cells (Ry9666) carrying *mCherry-TUB1* and *SCC1-yEGFP* fusions were analyzed by live imaging. **(A)** Representative violin plot showing metaphase-duration distribution for one *cdc13-1* single-cell dataset. Individual cells are plotted as points; boxplots indicate median and interquartile range. Metaphase-duration distributions were calculated among cells that entered anaphase. This analysis therefore describes the timing distribution of successful checkpoint bypass and is conditional on anaphase entry. **(B)** Representative time-lapse images illustrating the categories used to score post-mitotic rebudding after metaphase exit. Proliferative outcome was scored for the two compartments of the arrested mother-daughter pair: score 0, neither compartment rebudded; score 1, one compartment rebudded; and score 2, both compartments rebudded. **(C)** Representative single dataset analysis showing rebudding score as a function of metaphase duration. Dashed vertical lines mark the 350-min and 550-min descriptive reference boundaries. **(D-E)** Pooled analysis of four independent *cdc13-1* datasets (four biological replicates). Violin plots show metaphase-duration distributions (**D**), and scatter plots and violin show the relationship between metaphase duration and rebudding score (**E**). Individual points represent single-cells pooled from four independent datasets, with color gradients indicating dataset identity. Boxplots indicate the median and interquartile range. In the pooled population, score-0 cells had a median metaphase duration of 540 min (*n* = 81; IQR, 480-600 min), score-1 cells had a median of 450 min (*n* = 65; IQR, 390-495 min), and score-2 cells had a median of 390 min (*n* = 69; IQR, 345-435 min). **(F)** Logistic-regression model showing the probability of dual-compartment rebudding, corresponding to score 2, as a function of metaphase duration. **(G)** Classification and quantification of anaphase geometry. Cells were scored as undergoing symmetric or asymmetric anaphase, and anaphase type was compared across rebudding-score classes. Symmetric anaphase predominated and was enriched among cells with high rebudding scores. Data are shown as single-cell values and/or mean ± SEM, as indicated. **(H)** Metaphase duration according to anaphase geometry in *cdc13-1* cells. Violin plots show the distribution of metaphase duration among cells that entered anaphase, grouped by whether anaphase was scored as symmetric or asymmetric. Individual points represent single-cells pooled from four biological replicates (datasets), with color gradients indicating dataset identity. Boxplots indicate the median and interquartile range. Dashed horizontal lines mark the 350-min and 550-min descriptive reference boundaries. Statistical comparison was performed using a two-sided Mann-Whitney test. **p < 0.01.

We next asked whether the timing of checkpoint bypass predicted subsequent proliferative competence. Individual cells were followed from metaphase arrest through mitotic exit and subsequent cell-cycle re-entry, and the proliferative outcome of the arrested mother-daughter pair was scored according to whether the mother cell, the daughter cell, or both initiated a new cell cycle after adaptation. A score of 2 was assigned when both compartments rebudded, indicating complete proliferative re-entry; a score of 1 when only one compartment rebudded, indicating partial re-entry; and a score of 0 when neither compartment resumed proliferation (Fig. 2B). Cells that exited metaphase earlier most frequently achieved dual-compartment rebudding, whereas longer arrest was associated with a progressively lower probability of proliferation and an increased frequency of partial or failed rebudding (Fig. 2C). For descriptive visualization, we divided this distribution into early (<350 min), intermediate (350-550 min), and late (>550 min) arrest ranges. Full proliferative re-entry predominated in the early range, all three outcomes were observed in the intermediate range, and failed re-entry was most frequent in the late range. These boundaries were used as descriptive reference ranges rather than as thresholds for statistical inference (Fig. 2C). To test the reproducibility of this relationship, we analyzed four independent *cdc13-1* biological replicates, including one acquired on a different imaging platform. Absolute metaphase-duration distributions differed between experiments, and these differences were only partly affected by outlier removal (Supplementary Fig. 2A-C). Nevertheless, the association between timing and outcome was preserved (Supplementary Fig. 2D). Pooling the datasets reinforced a robust inverse relationship between metaphase duration and full proliferative capacity. Among cells that eventually bypassed the checkpoint, longer pre-bypass metaphase arrest was associated with a progressively lower probability of proliferative re-entry (Fig. 2D-E). We therefore analyzed metaphase duration as a continuous predictor of proliferative outcome across the four aggregated datasets. Full proliferative re-entry was defined as rebudding of both compartments of the arrested mother-daughter pair, whereas cells with rebudding of one or neither compartment were classified as showing incomplete proliferative capacity. After accounting for dataset-to-dataset variation, each additional 100 min of metaphase arrest markedly reduced the odds of complete proliferative re-entry (OR, 0.20; 95% CI, 0.117-0.341; P = 3.25 × 10^-9^; Fig. 2F and Supplementary Table 2). Analysis of the complete 0-1-2 rebudding score gave the same conclusion. Thus, metaphase duration predicts proliferative competence as a continuous variable, whereas the 350- and 550-min boundaries provide descriptive reference ranges for visualizing this relationship (Fig. 2E-F).

We next asked whether prolonged metaphase arrest was associated with altered anaphase geometry, which could reflect errors in spindle orientation or nuclear positioning. Following checkpoint bypass, cells underwent either symmetric anaphase, in which spindle elongation and chromosome segregation occurred across the mother-daughter neck, or asymmetric anaphase, in which segregation remained confined to one compartment (Fig. 2G). Asymmetric anaphases were associated with longer metaphase arrests, with a median duration of 510 min versus 450 min for symmetric divisions (Mann-Whitney U test, p = 0.0029; Fig. 2H). Although asymmetric divisions were never followed by full rebudding, symmetric divisions produced all rebudding outcomes, including complete failure. Thus, longer arrest was associated with an increased frequency of abnormal anaphase geometry, consistent with defects in spindle orientation or nuclear positioning, but these defects alone do not explain the loss of proliferative competence.

### Mad1-dependent SAC activity shifts arrest timing without abolishing the duration-outcome relationship

Previous studies using defined *HO*-induced double-strand breaks showed that spindle assembly checkpoint components help maintain prolonged DNA-damage-induced arrest, consistent with SAC activity arising from chromosome-attachment defects during persistent metaphase [40, 41]. We therefore asked whether Mad1-dependent SAC activity influences the timing of checkpoint bypass, the relationship between arrest duration and proliferative outcome, or both. For this analysis, we compared *cdc13-1* and *cdc13-1 mad1Δ* cells in a shared isogenic background, distinct from that used in the reference *cdc13-1* experiments (Supplementary Table 1). Metaphase duration was significantly shorter in *cdc13-1 mad1Δ* cells than in their isogenic *cdc13-1* controls, both before and after outlier removal (Fig. 3A and Supplementary Fig. 3A). Despite this shift, metaphase duration remained inversely associated with complete proliferative re-entry (Fig. 3B and Supplementary Fig. 3B). *MAD1* deletion did not substantially alter the overall frequency of anaphase entry and caused only a modest reduction in asymmetric divisions under these conditions (Fig. 3C). Thus, at 32°C, Mad1-dependent SAC activity prolongs the checkpoint-associated metaphase arrest but is not required for the inverse association between arrest duration and subsequent proliferative competence. When applied to the matched *cdc13-1* and *cdc13-1 mad1Δ* strains, the original timing model underestimated the absolute frequency of complete proliferative re-entry in both genotypes, indicating imperfect calibration across experimental backgrounds. Nevertheless, direct refitting within each strain confirmed that metaphase duration remained a significant negative predictor of complete proliferative re-entry (Fig. 3D). Thus, the duration–outcome relationship was reproduced across *cdc13-1* backgrounds, although the original model did not accurately predict absolute outcome probabilities in the matched strains. Because SAC activity might become particularly important when telomere dysfunction is more severe, we first examined the effect of increasing the temperature from 32°C to 34°C in the reference *cdc13-1* background. At 34°C, metaphase durations shifted toward longer values (Supplementary Fig. 3C), and complete proliferative re-entry was markedly reduced (Supplementary Fig. 3D). The model generated at 32°C continued to rank outcomes according to metaphase duration at 34°C but overestimated the absolute probability of complete proliferative re-entry. Models fitted directly to each condition retained an inverse association between metaphase duration and complete rebudding, with a lower predicted probability of complete proliferative re-entry at 34°C for comparable arrest durations (Supplementary Fig. 3E). These results indicate that the more restrictive telomere-uncapping condition altered both arrest timing and the probability of proliferative re-entry at a given arrest duration.

**Figure 3.**
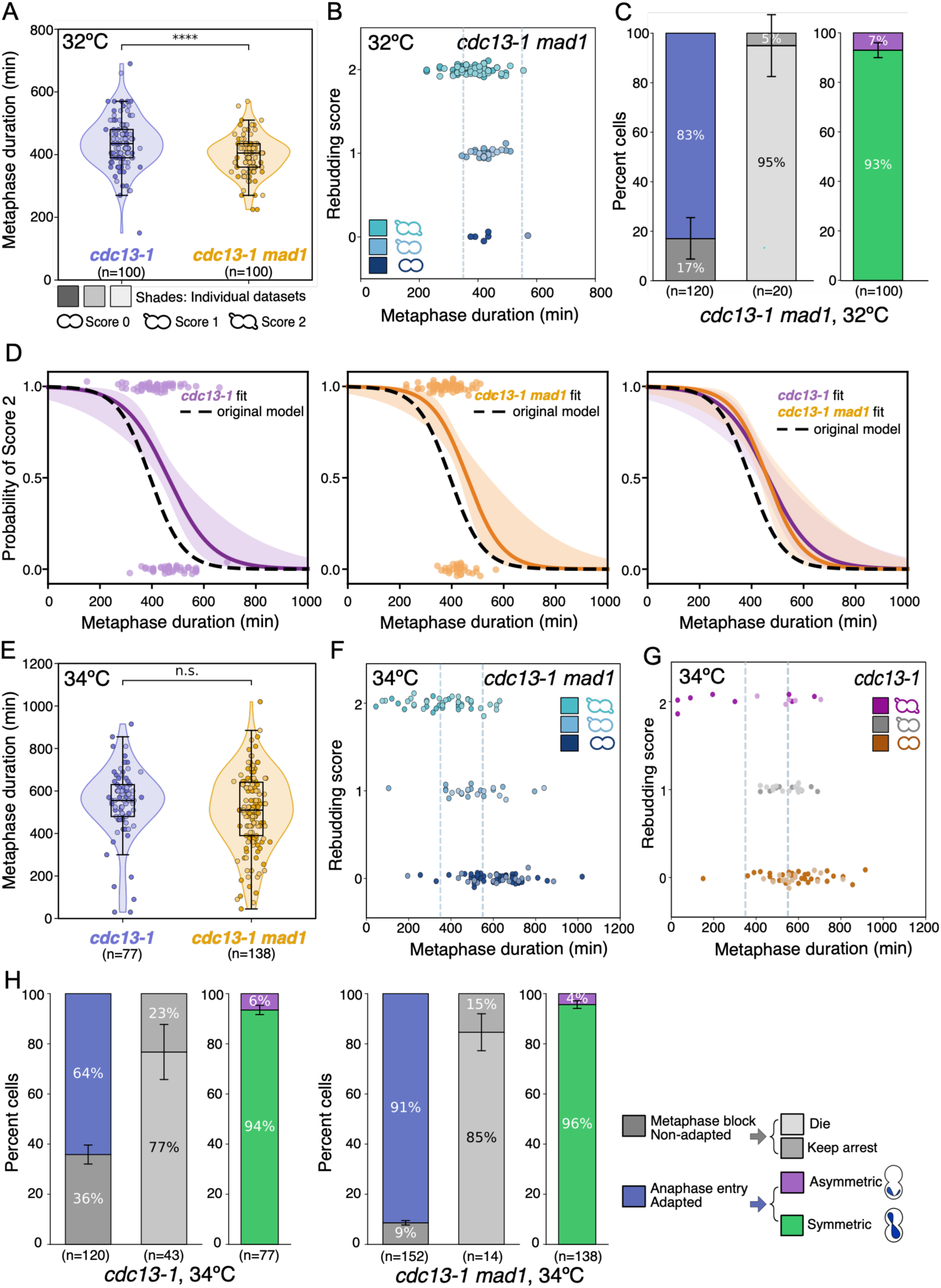
Loss of *MAD1* shifts arrest timing without abolishing the duration-outcome relationship. **(A)** Violin plots show the distribution of metaphase duration in individual *cdc13-1* (Ry6406) and *cdc13-1 mad1Δ* (Ry8843) cells carrying *HTB2-mCherry* and *TUB1-GFP* fusions at 32°C, with overlaid single-cell measurements from two independent pools per genotype. Boxplots indicate the median and interquartile range. The graph shows all cells, including outliers. Statistical comparisons were performed using two-sided Mann-Whitney tests. ****p < 0.0001. **(B)** Rebudding score plotted as a function of metaphase duration for *cdc13-1 mad1Δ* cells at 32°C. Each dot represents one cell. Cells are grouped by rebudding score (0, 1, or 2) and plotted according to metaphase duration. Different shades within each color indicate cells derived from independent experiments. Dashed vertical lines mark the 350 min and 550 min descriptive reference boundaries. **(C)** Quantification of adaptation outcome, fate of non-adapting cells, and anaphase geometry in *cdc13-1 mad1Δ* cells at 32°C. Stacked bars show the proportion of cells entering anaphase or remaining metaphase-arrested, the fate of metaphase-arrested cells, and the frequency of symmetric or asymmetric anaphase among adapted cells. Left, percentage of cells that either remained arrested in metaphase (Metaphase block/non-adapted, gray) or entered anaphase (Anaphase entry/adapted, blue). Center, fate of the non-adapting / metaphase-arrested cell fraction, classified as dead (light gray) or remained arrested (dark gray). Right, anaphase geometry of cells that entered anaphase, symmetric (green) versus asymmetric (purple). Bars show pooled percentages, and error bars indicate SEM calculated from two independent experiments. Because these data derive from two independent experiments, the graph reports pooled percentages with SEM, but no formal significance test was applied here. **(D)** Logistic-regression analysis showing the probability of full proliferative capacity as a function of metaphase duration for *cdc13-1*, *cdc13-1 mad1Δ*, and the combined dataset. Shaded regions indicate confidence intervals; dashed lines show the original model fit. **(E)** Violin plots showing metaphase duration in *cdc13-1* and *cdc13-1 mad1Δ* cells at 34°C. With all cells included, metaphase duration was not significantly different between genotypes (median 555 versus 510 min; Mann-Whitney U test, p = 0.0823). **(F-G)** Rebudding score plotted as a function of metaphase duration at 34°C in *cdc13-1 mad1Δ* (**F**) and *cdc13-1* (**G**) cells. **(H)** Quantification of adaptation outcome, fate of non-adapting cells, and anaphase outcome in *cdc13-1* and *cdc13-1 mad1Δ* cells at 34°C. Loss of *MAD1* significantly increased anaphase entry from 64.0 ± 3.8% to 91.3 ± 0.9% (Welch’s t-test, p = 0.0148), but did not significantly alter death among metaphase-arrested cells or the frequency of symmetric anaphase among adapted cells. Bars show pooled percentages; error bars indicate SEM from three independent datasets. Metaphase-fate percentages were calculated within the metaphase-arrested population, and anaphase-outcome percentages were calculated within the anaphase-entry population.

We next examined how SAC activity influenced adaptation at 34°C using the matched isogenic *cdc13-1* and *cdc13-1 mad1Δ* strains. At 34°C, *MAD1* deletion shifted metaphase duration toward shorter values, although this difference did not reach significance in the complete dataset (*P* = 0.0823) and was significant only in the outlier-exclusion sensitivity analysis (Fig. 3E and Supplementary Fig. 3F). In the isogenic *cdc13-1* control, most cells entering anaphase failed to achieve complete proliferative re-entry. By contrast, *cdc13-1 mad1Δ* cells displayed a broader range of outcomes: full proliferative re-entry was associated with shorter metaphase durations, partial proliferation with intermediate durations, and failure to rebud with the longest arrests (Fig. 3F-G and Supplementary Fig. 3G). Logistic modeling confirmed that metaphase duration remained a negative predictor of complete proliferative re-entry in both genotypes at 34°C.

After accounting for metaphase duration, *cdc13-1 mad1Δ* cells showed a higher probability of complete proliferative re-entry than matched *cdc13-1* controls, although this did not reach statistical significance (adjusted OR, 2.21; 95% CI, 0.96-5.06; *P* = 0.061). The duration x genotype interaction was also not significant (*P* = 0.124), providing no evidence that *MAD1* deletion altered the slope of the duration-outcome relationship (Supplementary Fig. 3H and Supplementary Table 2). *MAD1* deletion also increased anaphase entry from 64.0 ± 3.8% to 91.3 ± 0.9%, without significantly changing death among cells that remained arrested or the proportion of symmetric divisions among adapted cells (Fig. 3H). Together, these results indicate that Mad1-dependent SAC activity shapes the temporal range of checkpoint arrest and, under more severe telomere dysfunction, restrains checkpoint bypass. However, Mad1 is not required for the inverse association between arrest duration and post-bypass proliferative competence.

### Damage context is associated with distinct adaptation-time and outcome distributions

Having established the timing-outcome relationship after telomere uncapping, we asked whether a defined DNA double-strand break generated a similar distribution of arrest durations. We therefore compared *cdc13-1* cells, which experience persistent and heterogeneous telomere dysfunction, with *GAL-HO* cells in which a single DNA double-strand break was induced [4, 8, 9, 32].

Metaphase duration remained heterogeneous among *GAL-HO* cells but occupied a narrower range than in *cdc13-1* cells. *GAL-HO* durations were centered at approximately 420 min, whereas *cdc13-1* cells displayed a broader, right-skewed distribution extending into prolonged arrest; this difference persisted after exclusion of extreme values (Fig. 4A-B and Supplementary Fig. 4A-C). Complete proliferative re-entry predominated among the *GAL-HO* cells included in the duration–outcome analysis, with 222 of 249 cells achieving a rebudding score of 2. Within this narrower temporal range, in which incomplete or failed rebudding was uncommon, no significant association between metaphase duration and rebudding outcome was detected (Fig. 4C and Supplementary Fig. 4D).

**Figure 4.**
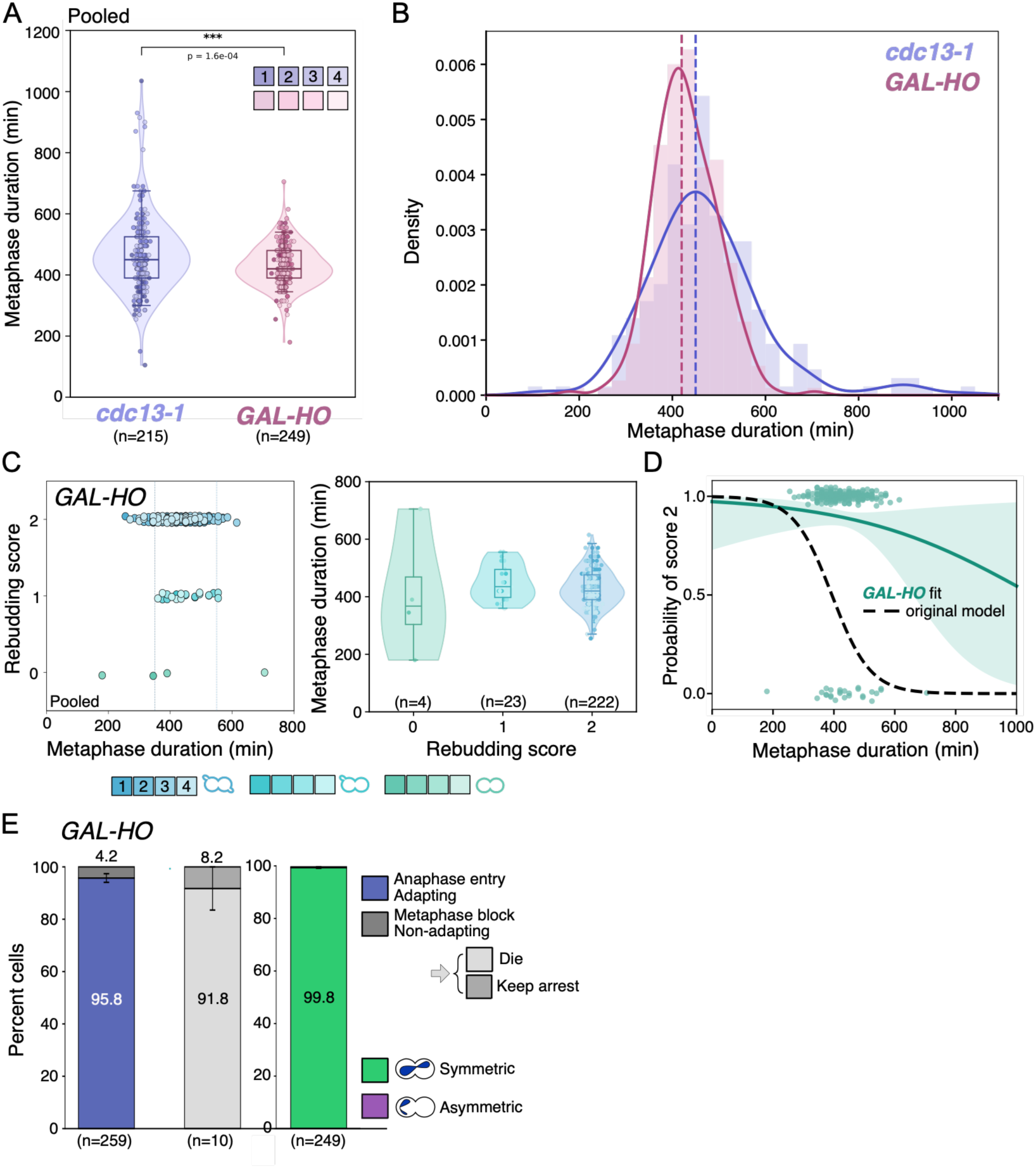
*GAL-HO* cells show a narrower metaphase-duration distribution and robust proliferative outcomes after DNA damage. **(A)** Violin plots comparing metaphase duration in pooled *cdc13-1* (Ry9666) and *GAL-HO* (Ry8847) single-cell datasets (4 datasets each). Individual points represent cells, and boxplots indicate median and interquartile range. *cdc13-1* cells showed significantly longer metaphase duration than *GAL-HO* cells (median 450 versus 420 min; Mann-Whitney U test, p = 1.60 × 10⁻⁴). **(B)** Density distribution of metaphase duration in *cdc13-1* and *GAL-HO* cells. Dashed lines indicate median values. *cdc13-1* cells display a broader, right-skewed distribution, whereas *GAL-HO* cells show a more compact distribution centered around intermediate durations. **(C)** Metaphase duration plotted as a function of rebudding score in *GAL-HO* cells. Each dot represents a single-cell; scores indicate no rebudding (score 0), intermediate rebudding (score 1), or robust rebudding (score 2). In the pooled population, score 0 comprised 4 cells, score 1 comprised 23 cells and score 2 comprised 222 cells. Median metaphase durations were 367.5 min for score 0 (IQR, 303.8-468.8 min), 435 min for score 1 (IQR, 397.5-495 min) and 420 min for score 2 (IQR, 390-476.3 min). Metaphase duration did not differ significantly among rebudding-score classes (Kruskal-Wallis test, *H* = 3.35, *P* = 0.187). Pairwise two-sided Mann-Whitney U tests with Holm correction were also not significant: score 0 versus score 1, adjusted *P* = 0.462; score 0 versus score 2, adjusted *P* = 0.462; and score 1 versus score 2, adjusted *P* = 0.429. No significant monotonic association was detected between metaphase duration and rebudding score (Spearman’s ρ = −0.056, *P* = 0.381). The 350- and 550-min reference boundaries were used for descriptive visualization and not for statistical inference. Violin plots summarize metaphase-duration distributions across rebudding-score classes. Most *GAL-HO* cells showed robust rebudding. **(D)** Logistic-regression analysis showing the probability of full proliferative capacity as a function of metaphase duration in *GAL-HO* cells, compared with the *cdc13-1* model. **(E)** Quantification of *GAL-HO* cell outcomes following *HO*-induced DNA damage. Stacked bars show adaptation outcome, fate of non-adapting/metaphase-arrested cells, and anaphase outcome among cells entering anaphase. Data are mean ± SEM from four independent datasets. No inferential statistical test was applied to the *GAL-HO* outcome bars because only one genotype/condition is shown.

Consistent with these differences in temporal and outcome range, the *cdc13-1* timing model showed limited discrimination and underestimated the frequency of complete proliferative re-entry in *GAL-HO* cells (Fig. 4D). *GAL-HO* cells rarely sampled the prolonged-arrest range in which proliferative failure was most evident after telomere uncapping, and most cells entered anaphase and underwent symmetric divisions (Fig. 4E). Thus, the two damage contexts occupy distinct adaptation-time and outcome ranges, and the timing-outcome relationship defined after telomere uncapping should not be interpreted as a universal clock for checkpoint adaptation.

### DNA-end metabolism alters arrest timing and baseline proliferative competence

The broad distribution of metaphase durations observed in *cdc13-1* cells suggested that DNA-end processing might influence both when cells bypass the checkpoint and their capacity to proliferate after bypass. We therefore analyzed mutants affecting distinct steps of DNA-end metabolism including *YKU70* [42–44], which limits resection at telomeres, the long-range resection factors *SGS1* and *EXO1* [45, 46], and the recombination regulator *SRS2* [4, 47]. *EXO1* was analyzed separately because its deletion delayed checkpoint establishment and prevented direct alignment with the synchronous first-cycle adaptation trajectories.

Among cells that successfully entered anaphase, *cdc13-1 yku70Δ, cdc13-1 sgs1Δ*, and *cdc13-1 srs2Δ* mutants all showed significantly shifted metaphase-duration distributions relative to *cdc13-1* (Fig. 5A). These differences persisted after exclusion of extreme values, indicating that DNA-end metabolism alters the timing distribution of successful checkpoint bypass rather than simply generating rare long-arrested cells (Supplementary Fig. 5A). Despite these shifts in bypass timing, the inverse relationship between metaphase duration and proliferative competence was retained in all three mutant backgrounds (Fig. 5B-C). Genotype-specific logistic models confirmed a significant decline in the probability of complete proliferative re-entry with increasing metaphase duration in *yku70Δ*, *sgs1Δ* and *srs2Δ* cells. The 350- and 550-min boundaries are shown in Fig. 5B only as descriptive reference ranges. The mutants differed, however, in their baseline proliferative outcomes. *cdc13-1 yku70Δ* and *cdc13-1 sgs1Δ* cells retained substantial proliferative capacity and high anaphase-entry frequencies, whereas *cdc13-1 srs2Δ* cells showed reduced checkpoint bypass and increased death during metaphase arrest. Among cells entering anaphase, symmetric divisions predominated in all three mutants (Fig. 5D). Thus, DNA-end metabolism influences two separable features of checkpoint adaptation: the timing distribution of successful checkpoint bypass and the overall frequency with which bypass results in productive proliferation. Despite these context-dependent shifts, longer metaphase duration remained associated with a lower probability of post-bypass proliferative competence across all three mutant backgrounds.

**Figure 5.**
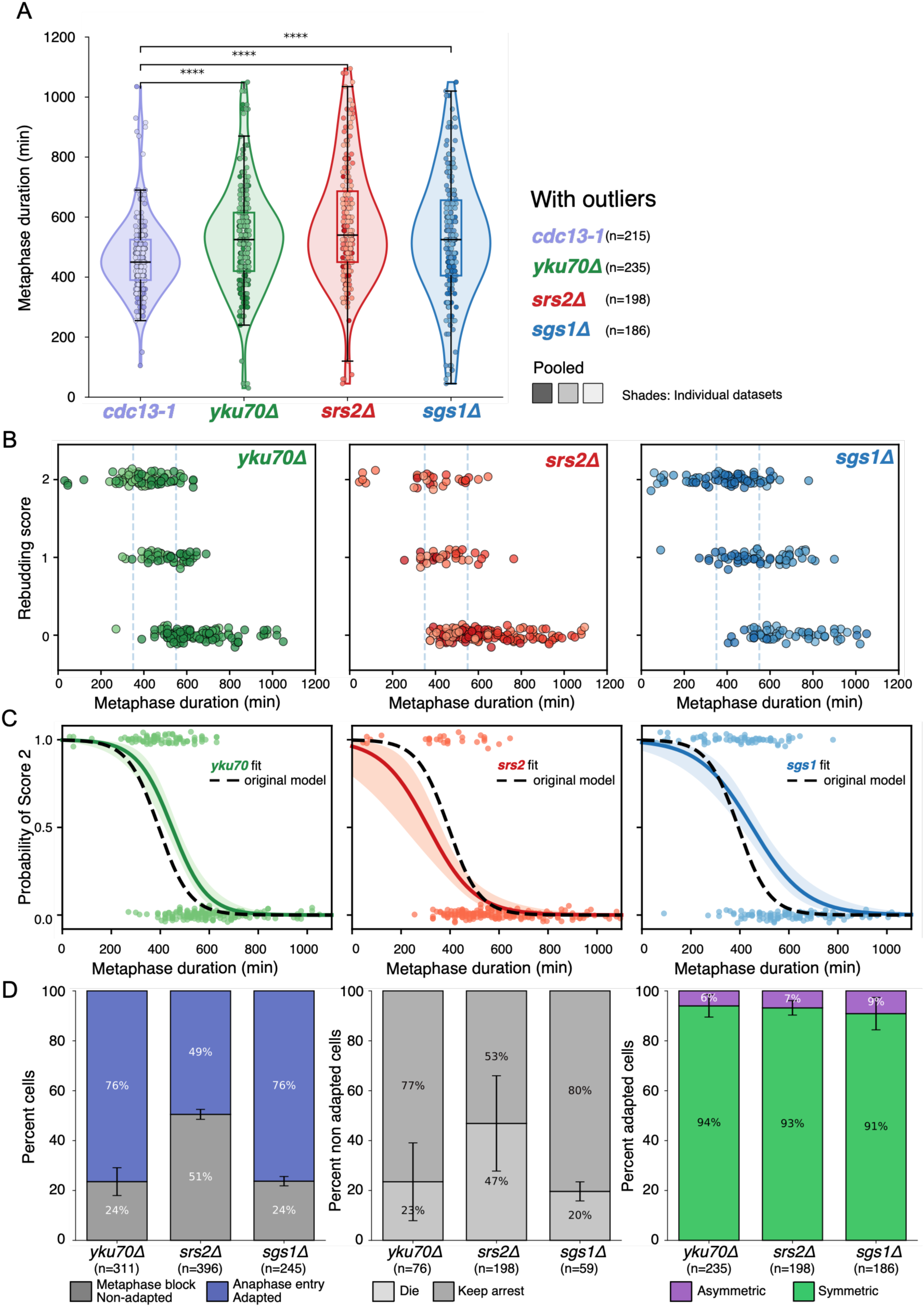
DNA-end metabolism mutants shift successful checkpoint-bypass timing and post-bypass proliferative outcome. **(A)** Violin plots showing metaphase-duration in *cdc13-1* (Ry9666), *cdc13-1 yku70Δ* (Ry11047), *cdc13-1 srs2Δ* (Ry11064), and *cdc13-1 sgs1Δ* (Ry10993) cells that entered anaphase. This analysis is conditional on anaphase entry and describes the timing distribution of successful checkpoint bypass. Individual points represent single-cells pooled from independent datasets, with point shade indicating dataset identity. Boxplots show the median and interquartile range. Median metaphase durations were 450 min for *cdc13-1*, 525 min for *yku70Δ*, 540 min for *srs2Δ*, and 525 min for *sgs1Δ*. Metaphase-duration distributions differed significantly across genotypes (Kruskal-Wallis test, p = 2.54 × 10⁻⁹). Pairwise two-sided Mann-Whitney U tests with Holm correction showed significant differences between *cdc13-1* and each mutant: *cdc13-1* versus *cdc13-1 yku70Δ*, adjusted *P* = 2.42 × 10⁻⁶; *cdc13-1* versus *cdc13-1 srs2Δ*, adjusted *P* = 1.63 × 10⁻⁹; and *cdc13-1* versus *cdc13-1 sgs1Δ*, adjusted *P* = 4.09 × 10⁻⁵. All measured cells, including statistical outliers, are shown; the corresponding analysis after genotype-specific outlier exclusion is presented in Supplementary Figure 5A. **(B)** Rebudding score as a function of metaphase duration among cells entering anaphase. Each dot represents a single-cell; dashed vertical lines indicate descriptive reference boundaries at 350 and 550 min. **(C)** Genotype-specific logistic-regression models showing the probability of dual-compartment rebudding as a function of metaphase duration. Colored curves indicate genotype-specific fits, shaded areas show confidence intervals, and the dashed black curve shows the original *cdc13-1* model for comparison. **(D)** Anaphase-entry frequency among all tracked cells, death among cells not entering anaphase, and asymmetric anaphase among cells with scorable anaphase geometry. *n* = cell number indicated in the figure.

### Checkpoint state determines how metaphase duration should be interpreted

We next used *rad9Δ* and *exo1Δ* cells to determine how checkpoint state affects the biological interpretation of metaphase duration and to distinguish checkpoint-associated arrest from the effects of persistent telomere damage itself. In *cdc13-1 rad9Δ* cells, Rad53 phosphorylation was not detected and cells progressed through mitosis without a sustained G2/M arrest (Supplementary Fig. 6A and Supplementary Video 3). These cells therefore did not undergo checkpoint adaptation in the strict sense, because they never established the arrested state from which adaptation occurs. Nevertheless, *rad9Δ* cells provided an informative boundary condition by separating telomere damage from checkpoint-mediated arrest. They continued to divide for multiple generations and retained high two-daughter output during early lineage events, indicating that damage *per se* is not sufficient to abolish proliferation acutely. Lineage output nevertheless declined at later events despite persistently short metaphases, showing that persistent or inherited telomere damage can progressively compromise proliferation independently of prolonged metaphase arrest (Supplementary Fig. 5B-D). In contrast to *cdc13-1 rad9Δ, cdc13-1 yku70Δ*, *cdc13-1 sgs1Δ* and *cdc13-1 srs2Δ* cells activated Rad53 with kinetics comparable to *cdc13-1*, confirming that they entered the canonical adaptation trajectory, although checkpoint persistence subsequently differed among the mutants (Supplementary Fig. 6B-D). These differences show that altered DNA-end metabolism can modify checkpoint persistence and mitotic timing after checkpoint establishment. *cdc13-1 exo1Δ* cells displayed a distinct boundary condition. Rad53 phosphorylation was delayed, and cells did not arrest synchronously during the first cell cycle but instead entered arrest after population synchrony had been lost (Supplementary Fig. 6E and Supplementary Video 4). Metaphase duration could therefore not be aligned to a common first-cycle checkpoint onset or compared directly with the other genotypes, and *exo1Δ* cells were excluded from the primary cross-genotype timing models. Nevertheless, single-cell lineage tracking allowed us to test whether the duration-outcome association remained detectable within the asynchronous *exo1Δ* trajectory (Supplementary Fig. 5B-D). For each mother lineage, we scored whether both expected daughters were produced in the subsequent event. In *exo1Δ* cells, metaphase duration increased substantially across successive lineage events. This progressive delay was accompanied by reduced two-daughter output.

In an analysis adjusted for lineage event, longer mother-cell metaphase duration was associated with a lower probability of producing both daughters. Thus, the duration-outcome association remained detectable in *exo1Δ* lineages, although metaphase duration could not be interpreted as the duration of a synchronously established checkpoint arrest. Instead, it captured progression through a delayed-checkpoint trajectory across successive mitoses (Supplementary Fig. 5E).

Together, these boundary conditions distinguish two contributions to declining proliferative output: persistence or inheritance of telomere damage across generations and increasing residence in a checkpoint-perturbed mitotic state. In *rad9Δ* cells, lineage output declined despite persistently short metaphases, demonstrating that damage can progressively compromise proliferation independently of prolonged checkpoint arrest. In *exo1Δ* cells, delayed checkpoint establishment was followed by progressive metaphase lengthening that was associated with reduced daughter output. Metaphase duration therefore reports different biological information depending on checkpoint state and should be viewed as a context-dependent trajectory variable rather than a universal proxy for damage burden.

## Discussion

Checkpoint adaptation and checkpoint recovery are conceptually distinct, but identifying either response requires separating checkpoint exit from its downstream consequences. Recovery describes resumption of cell-cycle progression after the initiating lesion has been repaired or reduced below the signaling threshold, whereas adaptation describes escape from an established checkpoint arrest despite persistent damage [4, 8, 9, 32]. Yet establishing that cells have adapted does not establish whether they retain the capacity to proliferate. Rad53 dephosphorylation, anaphase entry, rebudding and colony formation capture different aspects of this response, from checkpoint attenuation to subsequent proliferation, and should not be treated as interchangeable measures of adaptation [4, 9, 16, 35]. Our results make this distinction explicit: cells can enter anaphase after checkpoint arrest yet fail to regain complete proliferative competence. Among cells that bypassed the checkpoint, the duration of the preceding metaphase arrest predicted their subsequent proliferative outcome, linking the timing of checkpoint escape to its biological consequences.

The relationship between metaphase duration and proliferative outcome was continuous: among cells that entered anaphase, longer arrest was associated with a progressively lower probability of dual-compartment rebudding. This association was reproducible across independent experiments and remained detectable after perturbation of Mad1-dependent spindle checkpoint activity, increased telomere dysfunction and altered DNA-end metabolism. However, the absolute probability of complete proliferative re-entry at a given arrest duration varied across experimental backgrounds and conditions. Metaphase duration therefore provides predictive information within the telomere-dysfunction system, while the cellular context influences how that information translates into proliferative outcome. The 350- and 550-min boundaries illustrate regions of this continuous relationship rather than discrete biological thresholds. Metaphase duration should therefore not be interpreted as a discrete gate or universal molecular clock, but as a context-dependent variable reporting progression through a damaged and checkpoint-perturbed mitotic state.

The association between metaphase duration and proliferative outcome is predictive, but causality has not been established. Arrest duration and lesion state were not manipulated independently, and the relevant DNA structures were not measured in the same cells. Longer arrest could therefore contribute directly to declining proliferative competence, reflect progressive changes in the underlying lesion state, or integrate both processes. Metaphase duration should consequently be viewed as an integrated risk variable. It reports an increasing probability of post-bypass failure without identifying elapsed time itself as the cause.

The genetic perturbations further indicate that the timing of checkpoint bypass and the competence of cells to proliferate after bypass are related but separable features of adaptation. Loss of *MAD1* shifted the temporal range over which cells entered anaphase without abolishing the inverse association between metaphase duration and proliferative outcome. At 32°C, Mad1 loss shortened metaphase, whereas under more severe telomere dysfunction it also increased anaphase entry. Thus, Mad1-dependent spindle checkpoint activity influences how long cells remain arrested and, under stringent damage conditions, whether they undergo checkpoint bypass, but is not required for the duration-outcome association. DNA-end metabolism affected both bypass timing and proliferative outcome. Loss of *YKU70*, *SGS1* or *SRS2* shifted the distribution of successful checkpoint-bypass times, while the functional consequences differed among the mutants. *yku70Δ* and *sgs1Δ* cells retained substantial proliferative capacity, whereas *srs2Δ* cells combined delayed checkpoint bypass with reduced anaphase entry and increased death during arrest. Nevertheless, longer metaphase duration remained associated with poorer proliferative outcome in all three backgrounds. Together, these findings show that genetic perturbations can shift the temporal distribution of checkpoint bypass, the absolute probability of productive proliferation, or both, without eliminating the relationship between arrest duration and subsequent outcome.

The temporal and outcome landscape of adaptation also depended on damage context. Cells responding to a single *HO*-induced double-strand break occupied a comparatively narrow duration range in which complete proliferative re-entry predominated, whereas telomere uncapping generated a broader, right-skewed distribution extending into a prolonged-arrest, poor-outcome regime. The small number of *GAL-HO* cells that failed to rebud limited our ability to detect a duration– outcome association and should not be interpreted as evidence that timing is irrelevant after a double-strand break. Because the two experimental systems also differed in strain background, growth conditions and mode of damage induction, the observed distributions cannot be attributed exclusively to lesion architecture. Lesion number, chromosomal position and DNA-end structure may therefore contribute to the temporal range over which the consequences of checkpoint bypass become apparent [32, 40, 41].

The *rad9Δ* and *exo1Δ* boundary conditions further show that metaphase duration is not simply a proxy for the burden of telomere damage. *rad9Δ* cells did not establish a sustained checkpoint arrest and progressed through successive divisions with persistently short metaphases. Their high two-daughter output during early lineage events shows that telomere damage does not immediately abolish proliferation when checkpoint arrest is absent. Nevertheless, lineage output declined at later events, indicating that persistent or inherited damage can progressively compromise proliferation independently of prolonged metaphase arrest. By contrast, *exo1Δ* cells showed delayed checkpoint establishment: initial mitotic progression was followed by progressive metaphase lengthening and reduced two-daughter output across successive lineage events, with longer mother-cell metaphase duration remaining associated with poorer output after adjustment for lineage event. Together, these boundary conditions reveal two separable sources of proliferative vulnerability: persistence or inheritance of telomere damage across generations and increasing residence in a checkpoint-perturbed mitotic state. Checkpoint status therefore determines what metaphase duration reports; it should be viewed as a context-dependent trajectory variable rather than a universal proxy for damage burden.

What metaphase duration integrates at the molecular level remains unresolved. Prolonged arrest may accompany continued DNA-end processing, accumulation of recombination intermediates, persistent checkpoint exposure, altered mitotic-regulator dynamics, declining cellular homeostasis or defects in spindle function, nuclear positioning, chromosome segregation and mitotic exit. Anaphase geometry provides one measurable manifestation of this declining competence: longer arrests were associated with asymmetric divisions, and asymmetric anaphases were never followed by complete rebudding. However, symmetric divisions produced the full range of proliferative outcomes, indicating that gross segregation abnormalities contribute to failure but cannot fully explain the duration-outcome relationship. Cells with similar arrest durations may also carry different DNA structures and therefore differ in their ability to segregate chromosomes and resume proliferation.

The DNA-end-metabolism mutants provide additional clues to this heterogeneity. Srs2 may be particularly relevant because it acts at the interface between checkpoint disengagement and recombination control. By limiting RPA-associated checkpoint platforms and potentially restraining unresolved recombination intermediates, Srs2 may promote both checkpoint escape and preservation of a chromosome substrate compatible with mitosis [4, 47]. This interpretation remains a hypothesis, however, because the relevant DNA and recombination structures were not measured in the individual cells followed by live imaging. Recent work identifying spindle pole bodies as signaling platforms for Cdc5 during checkpoint adaptation [37], together with evidence that Cdc5 and Cdc14 coordinate Top2-dependent resolution of sister-chromatid linkages during mitosis [48], provides additional mechanistic context for how chromosome state at checkpoint bypass could influence productive segregation.

A further limitation is that checkpoint signaling was not monitored live in the individual cells used for fate and lineage analysis. We consequently cannot determine whether cells with similar metaphase durations experienced comparable checkpoint amplitudes or attenuation kinetics, nor can persistent population-level Rad53 phosphorylation be assigned to cells that subsequently divided or failed. This distinction is particularly relevant in *sgs1Δ* cells, in which substantial proliferation occurred despite persistent Rad53 phosphorylation at the population level. Combining a live checkpoint-activity reporter with Scc1 dynamics, spindle behavior and rebudding will be required to order checkpoint attenuation, anaphase entry and proliferative outcome within the same lineage. The FHA1-mCherry reporter described by Coutelier and colleagues provides one possible approach for monitoring Rad9-dependent signaling [49], although persistence of phosphorylated Rad9 after Rad53 dephosphorylation may limit its ability to report checkpoint attenuation directly.

The temporal organization identified here may be relevant beyond budding yeast. Human cells can escape sustained DNA-damage checkpoints and enter mitosis with persistent lesions, producing outcomes that include mitotic death, micronuclei formation and survival with damaged genomes [10, 11, 14, 28]. Prolonged mitosis can also be converted into post-mitotic arrest or death through mammalian mitotic-surveillance mechanisms [26, 27]. We do not infer that the timing relationship observed during yeast checkpoint adaptation represents the same molecular clock. Rather, our findings suggest the broader principle that the duration and quality of a damage-perturbed mitotic state may influence the consequences of checkpoint escape. This may be particularly relevant during telomere crisis, when dysfunctional chromosome ends generate fusions, anaphase bridges, micronuclei and chromothripsis [50, 51].

Together, our findings define DNA damage checkpoint adaptation as a temporally structured cell-fate trajectory rather than a binary arrest-versus-escape decision. Checkpoint bypass does not invariably restore proliferative competence: among cells entering anaphase after persistent damage, the duration of the preceding checkpoint-associated metaphase state predicts the probability of productive proliferation. Lesion context, checkpoint activity and DNA-end metabolism shape both the timing of bypass and the competence of cells to proliferate thereafter. Metaphase duration therefore functions as a context-dependent indicator of post-bypass outcome rather than a universal molecular clock. The cells with the greatest potential to propagate genome instability are consequently not simply those that divide with persistent damage, but those that do so while retaining the capacity for continued proliferation.

## Supplementary Data statement

Supplementary Data are available at *NAR* Online

## Acknowledgments

We thank M. Foiani, A. Rudner, A. Amon, A. Ciliberto and J.V. Kilmartin for strains and reagents; members of the R.V. laboratory for critical discussions and critical reading of the manuscript.

## Author contributions

A.D. and R.V. conceived and designed the study. A.D., E.C. and C.V. performed most of the original experimental work, with support from S.S. A.D., E.C., C.V. and R.V. analyzed the data; R.V. provided resources. R.V. wrote the manuscript and prepared the figures, with input from A.D., E.C., and C.V. All authors read and approved the manuscript.

## Data availability

All data supporting this study are available within the main text, figures and supplementary data file.

