## Supplementary material for "Timing of DNA damage checkpoint adaptation predicts post-mitotic proliferative competence": Dondi_Calabrese_Visintin_Supplementary_BioRxiv copy.pdf

##### This PDF file includes:

##### Six supplementary figures:

- **Supplementary Fig. 1.** Population-level and live-cell characterization of checkpoint signaling, mitotic progression and cell fate following telomere uncapping.
- **Supplementary Fig. 2**. Reproducibility of metaphase-duration measurements and their relationship with post-adaptation proliferative outcome in *cdc13-1* cells.
- **Supplementary Fig. 3**. Metaphase duration predicts post-adaptation proliferative outcome across SAC status and telomere-damage severity.
- **Supplementary Fig. 4**. Reproducibility of metaphase-duration measurements and their relationship with post-adaptation rebudding following a single HO-induced DNA double-strand break.
- **Supplementary Fig. 5.** Single-cell and lineage analyses distinguish DNA-end processing, checkpoint establishment and proliferative output.
- **Supplementary Fig. 6**. Checkpoint activation and cell-cycle progression in *cdc13-1* checkpoint and repair mutants after telomere uncapping.

##### Three supplementary Tables:

- **Supplementary Table 1:** Yeast strains
- **Supplementary Table 2.** Logistic-regression and model transfer analyses of metaphase duration and complete rebudding
- **Supplementary Table 3.** Dataset groups used for model training, model transfer and genotype or lesion-context comparisons

##### Four supplementary Videos:

- **Supplementary Video 1:** *cdc13-1* strain
- **Supplementary Video 2:** *cdc13-1 cdc5-ad* strain
- **Supplementary Video 3:** *cdc13-1 rad9Δ* strain
- **Supplementary Video 4:** *cdc13-1 exo1Δ* strain

##### One supplementary method:

- **Supplementary Method:** Predictive modelling, validation, cross-context transfer and refitting of the metaphase-duration outcome model

### Supplementary Figures

Dondi_Calabrese_Visintin Supplementary Figure 1

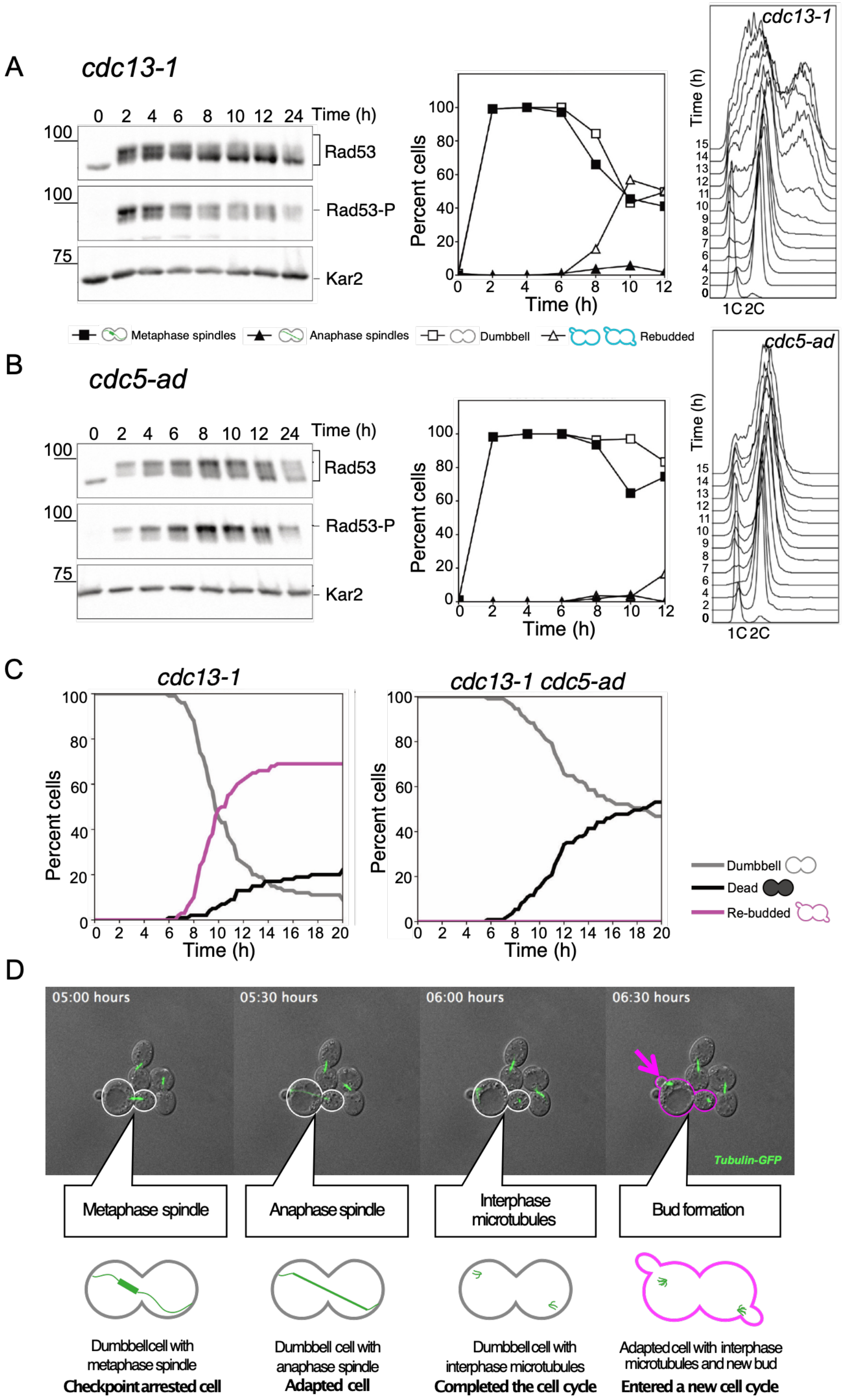

**Supplementary Fig. 1. Population-level and live-cell characterization of checkpoint signaling, mitotic progression and cell fate following telomere uncapping.**

**(A-B)** *cdc13-1* (Ry6406, **A**) and *cdc13-1 cdc5-ad* (Ry9161, **B**) cells carrying *HTB2-mCherry* and *TUB1-GFP* were synchronized in G1 and released at 32°C to induce telomere dysfunction. Samples were collected at the indicated times. Right panels show Rad53 abundance, electrophoretic mobility and phosphorylation by immunoblotting, with Kar2 used as a loading control. Middle panels show the percentages of cells displaying dumbbell morphology (open squares), rebudding (open triangles), metaphase spindles (closed squares) or anaphase spindles (closed triangles); $n$ = 100 cells per timepoint. Left panels show DNA-content profiles by flow cytometry. One representative experiment from three independent biological replicates per strain is shown.

**(C)** *cdc13-1* (Ry6090) and *cdc13-1 cdc5-ad* (Ry8525) cells were synchronized in G1, loaded into a microfluidic device and released from G1 arrest within the device at 32°C. Images were acquired every 15 min for 20 h. The percentages of cells displaying persistent dumbbell morphology, rebudding or cell death are shown over time; $n$ = 100 cells per strain.

**(D)** Representative time-lapse images of *cdc13-1* (Ry6406) cells carrying *HTB2-mCherry* and *TUB1-GFP,* released and imaged under the same microfluidic conditions as in (**C**), progressing from metaphase arrest through anaphase and subsequently rebudding. GFP-Tub1 is shown in green. Outlines indicate the tracked mother and daughter compartments before and after rebudding.

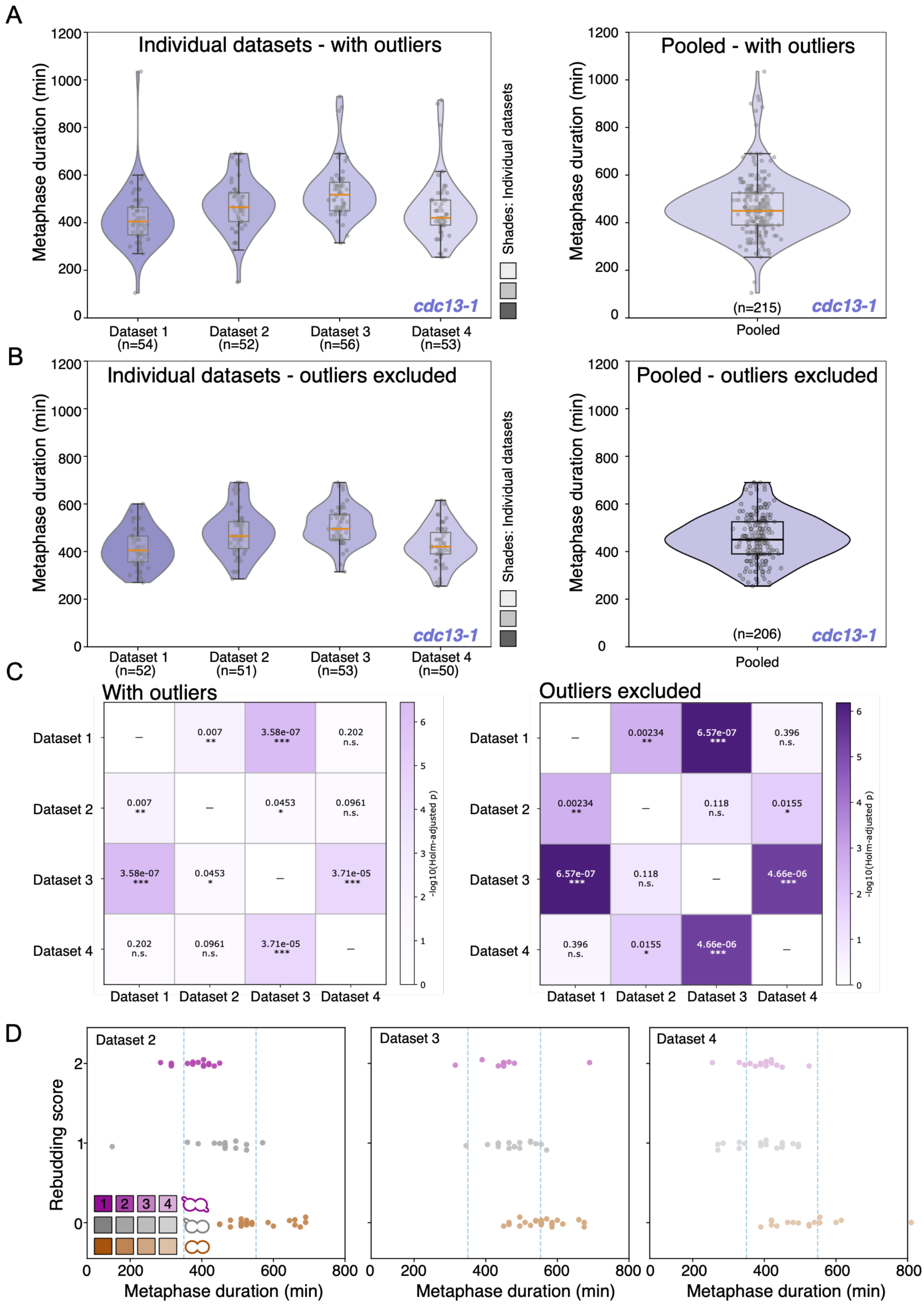


**Supplementary Fig. 2. Reproducibility of metaphase-duration measurements and their relationship with post-adaptation proliferative outcome in *cdc13-1* cells.**
Synchronously released *cdc13-1* cells (Ry9666) carrying *mCherry-TUB1 and SCC1-yEGFP* were analyzed by live-cell imaging in four independent experiments.

**(A)** Violin plots show the distribution of metaphase duration in each independent dataset and in the pooled population, with all measured cells included. Dataset 1 had a median metaphase duration of 405 min (IQR, 348.75-465 min), Dataset 2 of 465 min (IQR, 405-525 min), Dataset 3 of 517.5 min (IQR, 450-570 min) and Dataset 4 of 420 min (IQR, 390-495 min). Metaphase-duration distributions differed significantly among datasets (Kruskal-Wallis test, $H = 35.90$, $P = 7.87 \times 10^{-8}$).
**(B)** Metaphase-duration distributions after exclusion of outliers independently within each dataset using the 1.5 x interquartile-range criterion. Following outlier exclusion, the median metaphase durations were 405 min for Dataset 1 (IQR, 356.25-465 min), 465 min for Dataset 2 (IQR, 412.5-525 min), 495 min for Dataset 3 (IQR, 450-555 min) and 420 min for Dataset 4 (IQR, 390-480 min). In **(A, B)**, each point represents one cell; violins show the distribution density, boxes indicate the median and interquartile range, and whiskers extend to the most extreme values within 1.5 x the interquartile range.
**(C)** Pairwise comparisons of metaphase duration among the four datasets before and after outlier exclusion. Heatmaps report Holm-adjusted *P* values from two-sided Mann-Whitney U tests. Color intensity represents -log10 of the adjusted *P* value. n.s., not significant; *$P < 0.05$; **$P < 0.01$; ***$P < 0.001$; ****$P < 0.0001$. Outlier exclusion removed values of 105 and 1,035 min from Dataset 1; 150 min from Dataset 2; 870, 885 and 930 min from Dataset 3; and 810, 900 and 915 min from Dataset 4.
**(D)** Relationship between metaphase duration and post-adaptation rebudding outcome in three independent datasets. Each point represents one cell and is classified according to the rebudding score of the arrested mother-daughter pair: score 2, both compartments rebudded; score 1, only one compartment rebudded; and score 0, neither compartment rebudded. Vertical dashed lines indicate the descriptive temporal thresholds of 350 and 550 min.

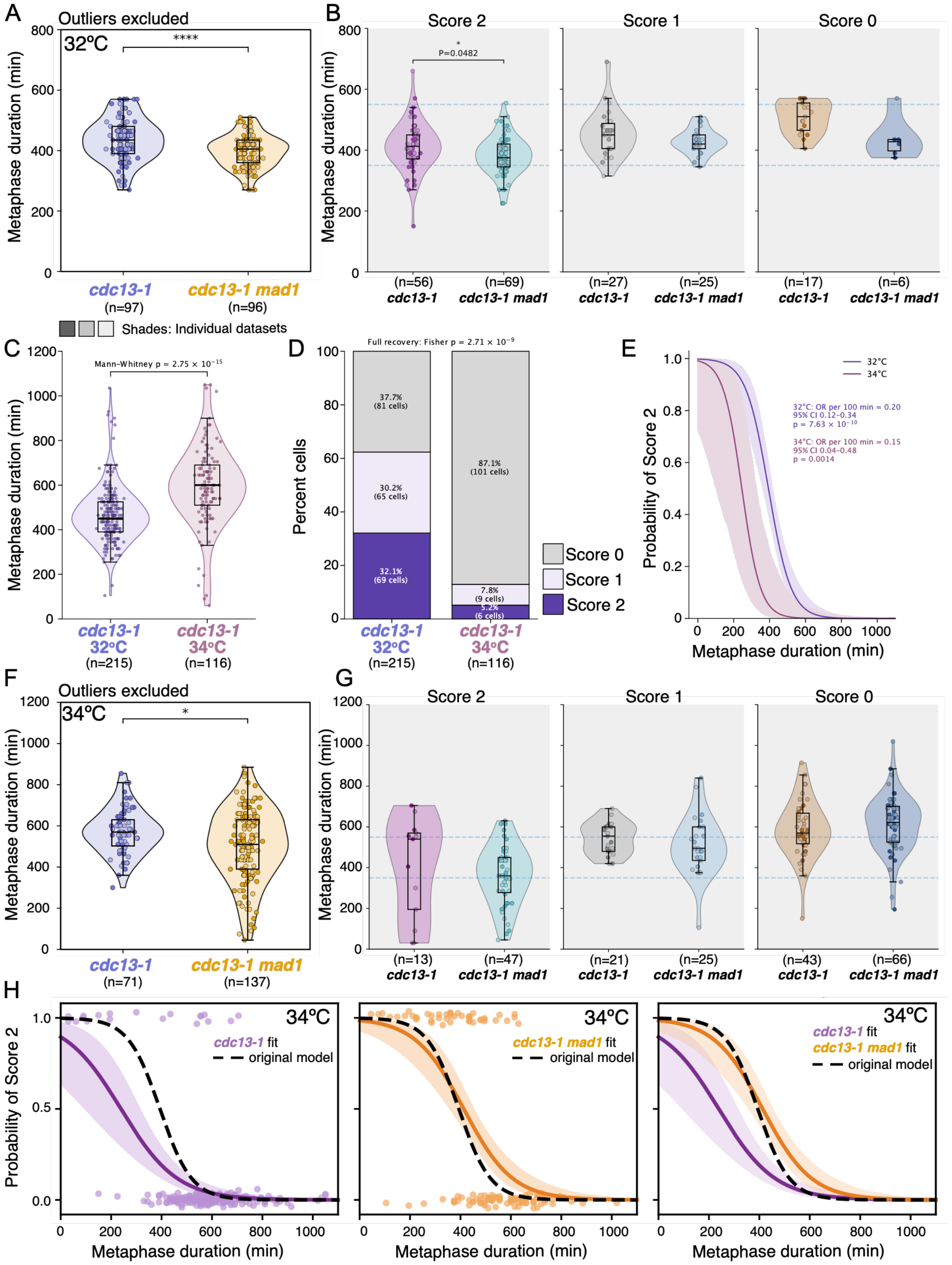


**Supplementary Fig. 3. Metaphase duration predicts post-adaptation proliferative outcome across SAC status and telomere-damage severity.**

**(A-B)** *cdc13-1* (Ry6406) and *cdc13-1 mad1Δ* (Ry8843) cells carrying HTB2-mCherry and TUB1-GFP were synchronized in G1 and released at 32°C to induce telomere dysfunction. Data from two independent datasets per genotype were pooled. **(A)** Violin plots show metaphase-duration distributions after exclusion of outliers using the 1.5 x interquartile-range criterion. Each point represents one cell; boxes indicate the median and interquartile range, and whiskers extend to the

most extreme values within 1.5 x the interquartile range. Metaphase-duration distributions were compared using a two-sided Mann-Whitney U test. ****$P$ < 0.0001. **(B)** Metaphase duration in cells that entered anaphase, stratified by post-mitotic rebudding outcome. A score of 2 indicates that both mother and daughter compartments rebudded, a score of 1 that only one compartment rebudded, and a score of 0 that neither compartment rebudded. Dashed horizontal lines indicate the descriptive temporal thresholds of 350 and 550 min. All cells with an observed metaphase-exit time and a scorable rebudding outcome were included in this analysis.
**(C-E)** *cdc13-1* cells (Ry9666) carrying mCherry-TUB1 and SCC1-yEGFP were synchronized in G1 and released at either 32°C or 34°C. **(C)** Violin plots show metaphase-duration distributions at the two temperatures. Each point represents one cell; boxes indicate the median and interquartile range, and whiskers extend to 1.5 x the interquartile range. **(D)** Distribution of post-mitotic proliferative outcomes at 32°C and 34°C. Bars show the number and percentage of cells assigned rebudding scores of 2, 1 or 0, as defined in **(B). (E)** Logistic-regression models showing the predicted probability of full proliferative re-entry, defined as a rebudding score of 2, as a function of metaphase duration at 32°C and 34°C. Cells with scores of 0 or 1 were classified as showing incomplete or failed proliferative re-entry. Shaded regions indicate 95% confidence intervals.
**(F-H)** *cdc13-1* (Ry6406) and *cdc13-1 mad1Δ* (Ry8843) cells carrying *HTB2-mCherry* and *TUB1-GFP* were synchronized in G1 and released at 34°C. **(F)** Violin plots show metaphase-duration distributions after exclusion of outliers using the 1.5 x interquartile-range criterion. **(G)** Metaphase duration in cells that entered anaphase, stratified by rebudding outcome as defined in (**B**). Dashed horizontal lines indicate the descriptive thresholds of 350 and 550 min. All cells with an observed metaphase-exit time and a scorable rebudding outcome were retained in this analysis. **(H)** Logistic-regression analysis of full proliferative re-entry at 34°C. Models fitted directly to the *cdc13-1* data (left) and *cdc13-1 mad1Δ* data (middle) are shown together with the original 32°C timing model generated in the reference *cdc13-1* background (black dashed line). The directly fitted curves for the two genotypes are overlaid in the right panel. Shaded regions indicate 95% confidence intervals. Unless otherwise indicated, each point represents one cell, and sample sizes are reported in the corresponding panels. Metaphase-duration distributions were compared using two-sided Mann-Whitney U tests, and rebudding-score distributions were compared using Fisher's exact test. Exact $P$ values are reported in the panels; *$P$ < 0.05 and ****$P$ < 0.0001. The 350- and 550-min thresholds were used for descriptive visualization and not for statistical inference.

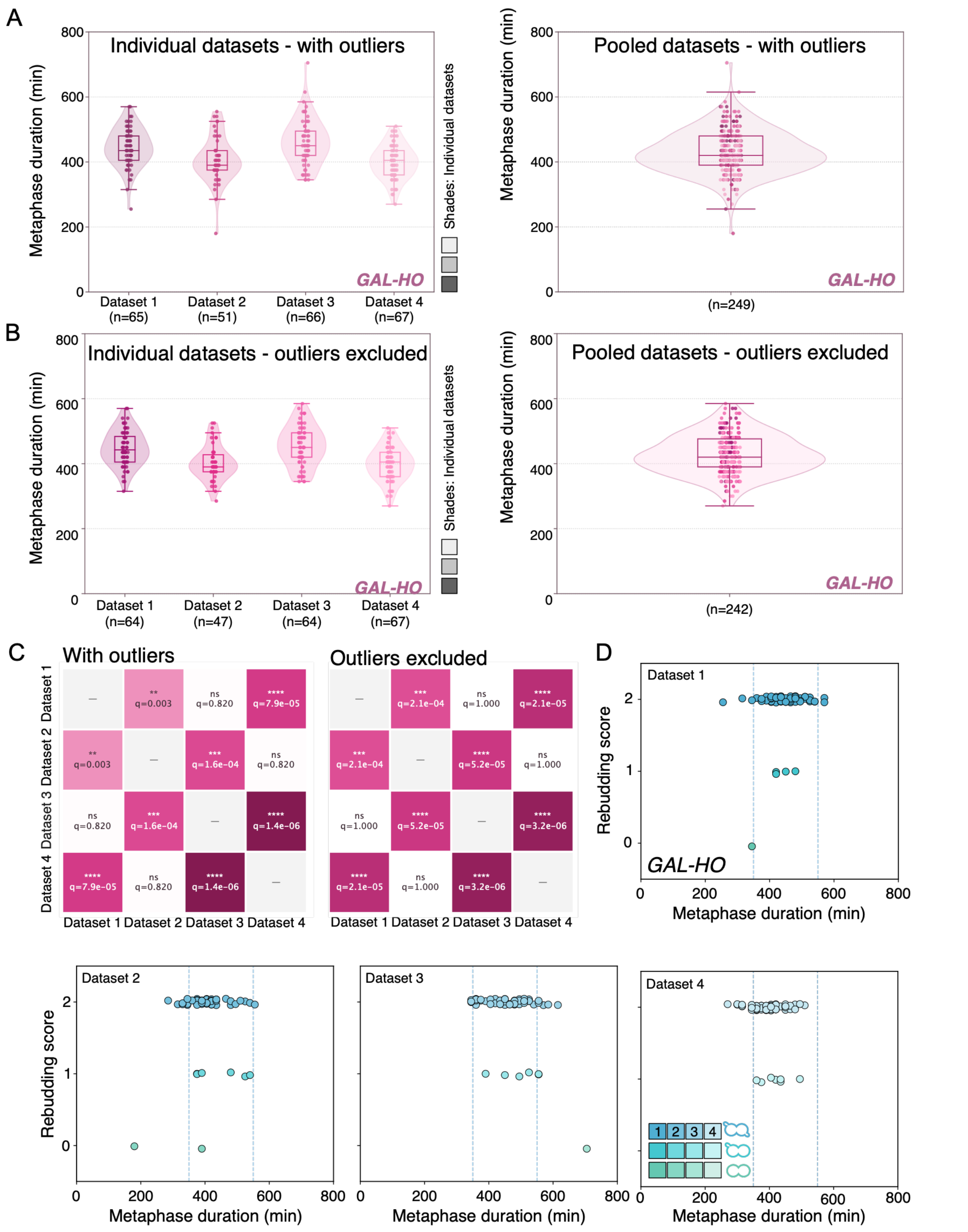


**Supplementary Fig. 4. Reproducibility of metaphase-duration measurements and their relationship with post-adaptation rebudding following a single HO-induced DNA double-strand break.**
*GAL-HO* cells (Ry8847) carrying *mCherry-TUB1* were synchronized in G1, loaded into a microfluidic device and released from G1 arrest within the device at 23°C into YEPR supplemented with 2% galactose to induce HO expression. Cells were analyzed by live-cell imaging in four independent

experiments. Metaphase duration was defined as the interval between formation of a short bipolar spindle and anaphase onset, identified by spindle elongation or disassembly.
**(A)** Violin plots show the distribution of metaphase duration in each independent dataset and in the pooled population, with all measured cells included. Dataset 1 had a median metaphase duration of 435 min (IQR, 405-480 min; $n$ = 65), Dataset 2 of 390 min (IQR, 375-435 min; $n$ = 51), Dataset 3 of 450 min (IQR, 420-495 min; $n$ = 66) and Dataset 4 of 405 min (IQR, 360-435 min; $n$ = 67). The pooled population had a median metaphase duration of 420 min (IQR, 390-480 min; $n$ = 249). Metaphase-duration distributions differed significantly among the four independent datasets (Kruskal-Wallis test, $H$ = 37.80, $P = 3.12 \times 10^{-8}$).
**(B)** Metaphase-duration distributions after exclusion of outliers independently within each dataset using Tukey's 1.5 x interquartile-range criterion. One value was excluded from Dataset 1, four from Dataset 2, two from Dataset 3 and none from Dataset 4. Following outlier exclusion, Dataset 1 had a median metaphase duration of 442.5 min (IQR, 405-483.8 min; $n$ = 64), Dataset 2 of 390 min (IQR, 375-427.5 min; $n$ = 47), Dataset 3 of 450 min (IQR, 420-495 min; $n$ = 64) and Dataset 4 of 405 min (IQR, 360-435 min; $n$ = 67). The pooled filtered population had a median metaphase duration of 420 min (IQR, 390-476.3 min; $n$ = 242). Metaphase-duration distributions remained significantly different after outlier exclusion (Kruskal-Wallis test, $H$ = 41.16, $P = 6.04 \times 10^{-9}$). In **(A, B),** each point represents one cell. Violins show the distribution density, and overlaid boxplots indicate the median and interquartile range; whiskers and caps extend to the most extreme values within 1.5 x the interquartile range. In the pooled plots, measurements from the four datasets were combined into a single distribution, while individual points retained their dataset-specific colors. The pooled population was not included as an independent group in statistical comparisons because it contains the same observations as the four individual datasets.
**(C)** Pairwise comparisons of metaphase duration among the four independent datasets before and after outlier exclusion. Heatmaps report Holm-adjusted $P$ values from two-sided Mann-Whitney U tests. Color intensity represents -log10 of the adjusted $P$ value. Exact adjusted $P$ values are reported in the heatmaps. n.s., not significant; *$P < 0.05$; **$P < 0.01$; ***$P < 0.001$; ****$P < 0.0001$.
**(D)** Relationship between metaphase duration and post-adaptation rebudding outcome in the four independent datasets pooled in Fig. 4C. Each point represents one cell. Rebudding outcome was scored for the arrested mother-daughter pair as follows: score 2, both compartments rebudded; score 1, only one compartment rebudded; and score 0, neither compartment rebudded. Colors denote rebudding-score classes, with dataset-specific shades used to distinguish the four independent experiments. Vertical dashed lines indicate the descriptive temporal thresholds of 350 and 550 min.

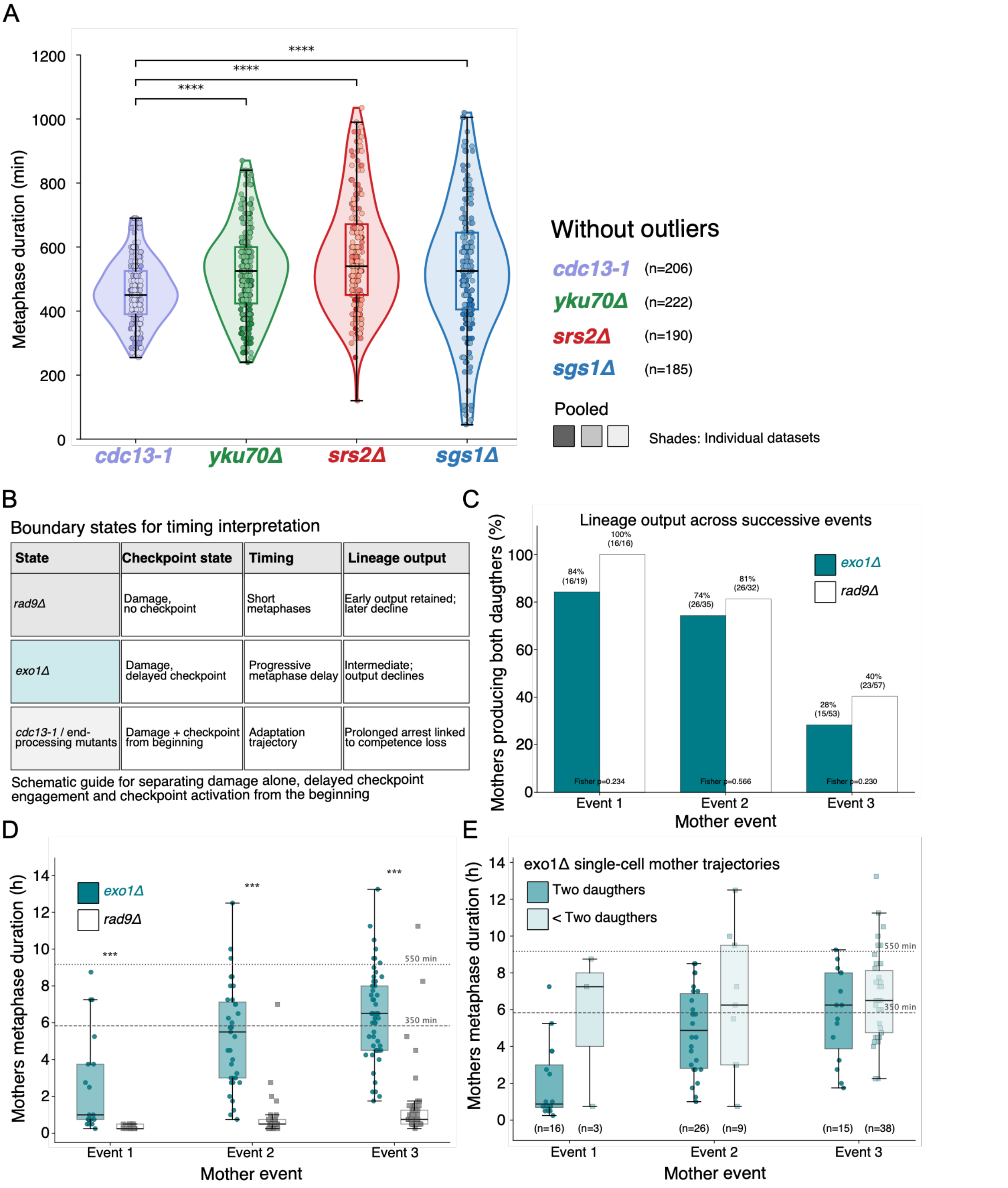


**Supplementary Fig. 5. Single-cell and lineage analyses distinguish DNA-end processing, checkpoint establishment and proliferative output.**
*cdc13-1* (Ry9666), *cdc13-1 yku70Δ* (Ry11047), *cdc13-1 srs2Δ* (Ry11064), *cdc13-1 sgs1Δ* (Ry10993), *cdc13-1 rad9Δ* (Ry10990) and *cdc13-1 exo1Δ* (Ry10998) cells carrying *mCherry-TUB1* and *SCC1-yEGFP* were synchronized in G1, loaded into a microfluidic device and released from G1 arrest within the device at 32°C to induce telomere dysfunction. Cells were followed by live-cell imaging. Metaphase duration was defined as the interval between formation of a short bipolar spindle and anaphase onset, identified by spindle elongation or disassembly together with loss of the Scc1 signal. (**A**) Violin plots show metaphase-duration distributions in *cdc13-1*, *cdc13-1 yku70Δ*, *cdc13-1 srs2Δ* and *cdc13-1 sgs1Δ* cells after exclusion of outliers independently within each genotype using Tukey's 1.5 x interquartile-range criterion. Data were pooled from four independent datasets. Each point represents one cell; boxes indicate the median and interquartile range, and whiskers extend to the most extreme values within 1.5 x the interquartile range. Metaphase-duration distributions differed

significantly among genotypes after outlier exclusion (Kruskal-Wallis test, $P = 1.82 \times 10^{-11}$). Pairwise comparisons were restricted to each mutant versus *cdc13-1* and adjusted across these three comparisons using Holm's method. Significant differences were detected between *cdc13-1* and *cdc13-1 yku70Δ* (adjusted $P = 3.11 \times 10^{-7}$), *cdc13-1* and *cdc13-1 srs2Δ* (adjusted $P = 1.02 \times 10^{-11}$), and *cdc13-1* and *cdc13-1 sgs1Δ* (adjusted $P = 5.78 \times 10^{-6}$).
**(B)** Conceptual summary of the checkpoint states used to define the scope of the timing analysis. *cdc13-1 rad9Δ* represents telomere damage in the absence of checkpoint establishment, whereas *cdc13-1 exo1Δ* represents delayed and progressively emerging checkpoint engagement. By contrast, *cdc13-1*, *cdc13-1 yku70Δ*, *cdc13-1 sgs1Δ* and *cdc13-1 srs2Δ* cells establish checkpoint signaling from the beginning of the observed trajectory and can therefore be analyzed within the canonical adaptation-timing framework**.**
**(C)** Lineage output across successive mother-cell events in *cdc13-1 exo1Δ* and *cdc13-1 rad9Δ* cells. For each mother-cell event, daughter output was scored as complete only when both expected daughters, X.1 and X.2, were detected in the subsequent lineage event. Bars show the percentage of mother-cell events producing both daughters after duplicated sheets were removed; numbers indicate n/N. Counts are shown above the bars. The two genotypes did not differ significantly at any event by two-sided Fisher's exact test: event 1, 16/19 versus 16/16, $P = 0.234$; event 2, 26/35 versus 26/32, $P = 0.566$; and event 3, 15/53 versus 23/57, $P = 0.230$.
**(D)** Mother-cell metaphase duration across successive lineage events in *cdc13-1 exo1Δ* and *cdc13-1 rad9Δ* cells. Each point represents one mother-cell event; boxes indicate the median and interquartile range, and whiskers extend to the most extreme values within 1.5 x the interquartile range. Dashed horizontal lines indicate the descriptive temporal thresholds of 350 and 550 min. Metaphase duration was significantly longer in *cdc13-1 exo1Δ* than in *cdc13-1 rad9Δ* cells at each event by two-sided Mann-Whitney U test: event 1, median 60 versus 15 min, $P = 5.43 \times 10^{-6}$; event 2, median 330 versus 30 min, $P = 8.26 \times 10^{-11}$; and event 3, median 390 versus 45 min, $P = 1.04 \times 10^{-16}$.
**(E)** Mother-cell metaphase duration in *cdc13-1 exo1Δ* lineages stratified according to subsequent daughter output. Mother-cell events were classified according to whether both expected daughters were produced or whether one or both daughters were not detected. Each point represents one mother-cell event; boxes indicate the median and interquartile range, and whiskers extend to the most extreme values within 1.5 x the interquartile range. Sample sizes are shown below each group, and dashed horizontal lines indicate the descriptive thresholds of 350 and 550 min. In a logistic-regression model including lineage event as a covariate, longer mother-cell metaphase duration was associated with a lower probability of producing both daughters. Each additional hour in metaphase was associated with an odds ratio of 0.80 for complete daughter output (95% CI, 0.67-0.96; $P = 0.0144$). Outlier exclusion was applied only to the metaphase-duration distributions shown in **(A);** all eligible and scorable lineage events were retained in the analyses shown in **(C-E).**

Dondi_Calabrese_Visintin_Supplementary Figure 6

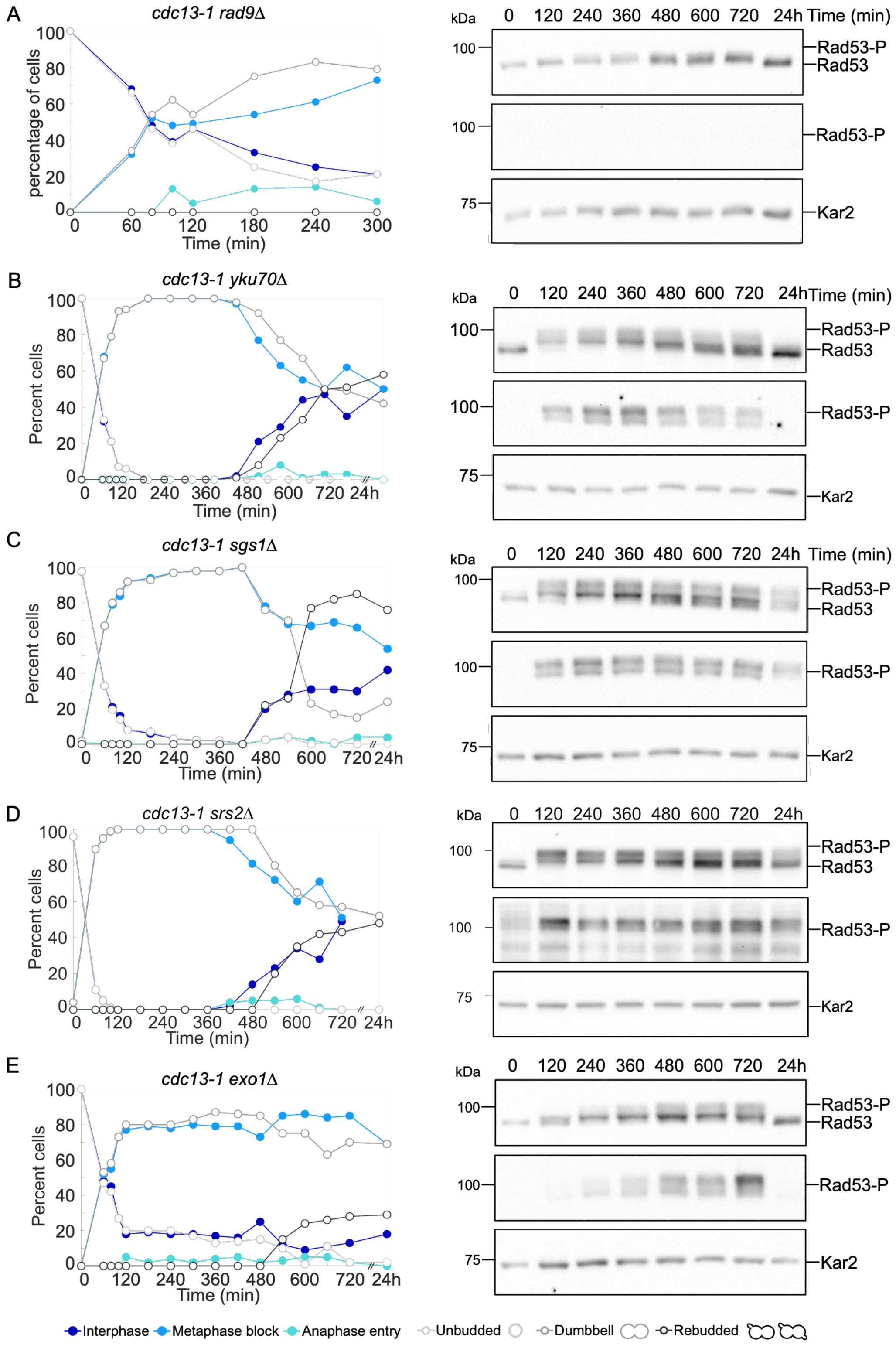


**Supplementary Fig. 6. Checkpoint activation and cell-cycle progression in *cdc13-1* checkpoint and repair mutants after telomere uncapping.**
**(A-E)** Bulk time-course analysis of *cdc13-1 rad9Δ* (Ry10990, **A**), *cdc13-1 yku70Δ* (Ry11047, **B**), *cdc13-1 sgs1Δ* (Ry10993, **C**), *cdc13-1 srs2Δ* (Ry11064, **D**), and *cdc13-1 exo1Δ* (Ry10998, **E**) cells carrying *mCherry-TUB1* and *SCC1-yEGFP* after release under telomere-uncapping conditions. One

representative experiment from three independent biological replicates *per* strain is shown. Right panels show immunoblots monitoring Rad53 phosphorylation as a marker of DNA damage checkpoint activation; Kar2 was used as a loading control. Time points are indicated in minutes after release; 24 h indicates the final time point. Left panels show the percentage of cells in each cell-cycle or fate category over time, including interphase, metaphase, anaphase, unbudded/binucleate, rebudded, and post-mitotic arrested cells, as indicated. *n*=100 cells *per* timepoint.

### Supplementary Tables

#### Supplementary Table 1: Yeast strains used in this study

| Name | Relevant Genotype | Origin |
|---|---|---|
| **W303** | *MATa, ade2-1, leu2-3, ura3, trp1-1, his3-11,15, can1-100, GAL, psi+, rad5-* | A. Amon |
| **SY2080** | *MATa, ade2-1, leu2-3, ura3, trp1-1, his3-11,15, can1-100, GAL, psi+, RAD5* | M. Foiani |
| **JKM179** | *MATα, hoΔ hmlΔ::ADE1 hmrΔ::ADE1, ade1-100, leu2-3,112, trp::hisG, lys5 ura3-52, ade3::GAL::HO* | J.E. Haber |
| **Ry6090** | *MATa, cdc13-1, RAD5+* | This study |
| **Ry8525** | *MATa, cdc13-1, cdc5-ad, RAD5+* | This study |
| **Ry6406** | *MATa, cdc13-1, HTB2-Cherry::HIS3, ura3::pAFS125-TUB1p-GFPTUB1::URA3, RAD5+* | This study |
| **Ry8843** | *MATa, cdc13-1, HTB2-Cherry::HIS3, ura3::pAFS125-TUB1p-GFPTUB1::URA3, mad1::hphMX6, RAD5+* | This study |
| **Ry8847** | *MATa, hoΔ hmlΔ::ADE1 hmrΔ::ADE1, ade1-100, leu2-3,112, trp1::hisG, lys5 ura3-52, ade3::GAL::HO, ura3::pRS306-mCherry-TUB1::URA3* | This study |
| **Ry9161** | *MATa, cdc13-1, cdc5-ad, HTB2-mCherry::HIS3, ura3::pAFS125-TUB1p-GFPTUB1::URA3, RAD5+* | This study |
| **Ry9666** | *MATa, cdc13-1, ura3::pRS306-mCherry-TUB1::URA3, SCC1-yEGFP::hphMX6, RAD5+* | This study |
| **Ry10642** | *MATa, cdc13-1, cdc5-ad, SCC1-yEGFP::hphMX6, ura3::pRS306-mcherry-TUB1::URA3, RAD5+* | This study |
| **Ry10990** | *MATa, cdc13-1, SCC1-yEGFP::hphMX6, ura3::pRS306-mcherry-TUB1::URA3, rad9::LEU2, RAD5+* | This study |
| **Ry10993** | *MATa, cdc13-1, SCC1-yEGFP::hphMX6, ura3::pRS306-mcherry-TUB1::URA3, sgs1::KanMX6, RAD5+* | This study |
| **Ry10998** | *MATa, cdc13-1, SCC1-yEGFP::hphMX6, ura3::pRS306-mcherry-TUB1::URA3, exo1::HIS3, RAD5+* | This study |
| **Ry11047** | *MATa, cdc13-1, SCC1-yEGFP::hphMX6, ura3::pRS306-mcherry-TUB1::URA3, yku70::HIS3, RAD5+* | This study |
| **Ry11064** | *MATa, cdc13-1, SCC1-yEGFP::hphMX6, ura3::pRS306-mCherry-TUB1::URA3, srs2::KanMX6, RAD5+* | This study |

#### Supplementary Table 2. Logistic-regression and model transfer analyses of metaphase duration and complete rebudding

| Analysis set | n | Observed complete rebudding | Expected from original model | Original-model AUC | Model / effect represented | OR | 95% CI | P value | Interpretation |
|---|---|---|---|---|---|---|---|---|---|
| **A. Primary timing model and sensitivity analyses** | | | | | | | | | |
| Original *cdc13-1* (Ry9666), 32°C: primary binary model | 215 | 69/215 | - | 0.849 | Complete rebudding ~ duration per 100 min + dataset | 0.20 | 0.117-0.341 | $3.25 \times 10^{-9}$ | Longer metaphase duration strongly reduced the probability of complete rebudding. |
| Original *cdc13-1* (Ry9666), 32°C: ordinal model | 215 | Scores 0/1/2 | - | - | Ordinal rebudding score ~ duration per 100 min + dataset | 0.16 | 0.10-0.25 | $1.58 \times 10^{-15}$ | Longer metaphase duration shifted cells toward lower rebudding scores. |
| Original *cdc13-1* (Ry9666), 32°C: outlier-exclusion sensitivity analysis | 206 | - | - | - | Complete rebudding ~ duration per 100 min + dataset after within-dataset 1.5 x IQR exclusion | 0.16 | 0.09-0.29 | $7.63 \times 10^{-10}$ | The duration effect was not driven by extreme long-arrest values. |
| **B. Matched *cdc13-1* and *cdc13-1 mad1Δ* cohorts at 32°C** | | | | | | | | | |
| Matched *cdc13-1* control (Ry6406), 32°C | 100 | 56/100 | 36.5/100 | 0.737 | Direct refit: duration per 100 min | 0.31 | 0.16-0.59 | $3.80 \times 10^{-4}$ | Duration remained predictive; the original model underestimated absolute complete rebudding. |
| Matched *cdc13-1 mad1Δ* (Ry8843), 32°C | 100 | 69/100 | 48.6/100 | 0.743 | Direct refit: duration per 100 min | 0.23 | 0.09-0.55 | $9.78 \times 10^{-4}$ | Duration remained predictive; the original model underestimated absolute complete rebudding. |
| Combined matched cohorts, 32°C | 200 | - | - | - | Duration effect in model: complete rebudding ~ duration + genotype | 0.27 | 0.16-0.46 | $1.52 \times 10^{-6}$ | Duration remained predictive across the two matched genotypes. |
| Combined matched cohorts, 32°C | 200 | - | - | - | *mad1Δ* versus *cdc13-1*, adjusted for duration | 1.08 | 0.57-2.07 | 0.814 | No duration-adjusted *MAD1* genotype effect was detected. |
| Combined matched cohorts, 32°C | 200 | - | - | - | Duration x *mad1Δ* interaction, adjusted for AD104/AD105 block | 0.72 | 0.24-2.17 | 0.560 | No evidence that *MAD1* deletion changed the duration-outcome slope. |
| **C. Increased telomere-damage severity and matched *MAD1* analysis at 34°C** | | | | | | | | | |
| Original *cdc13-1* (Ry9666), 34°C | 116 | 6/116 | 16.7/116 | 0.987 | Direct duration-only refit, per 100 min | 0.15 | 0.045-0.48 | 0.0014 | The original model overestimated absolute complete rebudding but strongly ranked outcomes; duration remained predictive. |
| Matched *cdc13-1* control (Ry6406), 34°C | 77 | 13/77 | 13.5/77 | 0.687 | Direct refit: duration per 100 min + dataset | 0.53 | 0.35-0.81 | 0.0032 | Duration remained predictive in the matched control background. |
| Matched *cdc13-1 mad1Δ* (Ry8843), 34°C | 138 | 47/138 | 38.9/138 | 0.869 | Direct refit: duration per 100 min + dataset | 0.34 | 0.23-0.50 | $4.12 \times 10^{-8}$ | Duration remained predictive after *MAD1* deletion. |
| Combined matched cohorts, 34°C | 215 | - | - | - | Duration effect in model: complete rebudding ~ duration + genotype | 0.42 | 0.32-0.55 | $2.82 \times 10^{-10}$ | Duration remained strongly predictive across the matched genotypes. |
| Combined matched cohorts, 34°C | 215 | - | - | - | *mad1Δ* versus *cdc13-1*, adjusted for duration | 2.21 | 0.96-5.06 | 0.061 | *MAD1* deletion showed a trend toward higher complete rebudding after accounting for duration. |
| Combined matched cohorts, 34°C | 215 | - | - | - | Duration x *mad1Δ* interaction | 0.65 | 0.38-1.12 | 0.124 | No evidence that *MAD1* deletion changed the duration-outcome slope at 34°C. |
| **Analysis set** | **n** | **Observed complete rebudding** | **Expected from original model** | **Original-model AUC** | **Model / effect represented** | **OR** | **95% CI** | **P value** | **Interpretation** |
| **D. Lesion-context and DNA-end-processing analyses** | | | | | | | | | |
| *GAL-HO* | 249 | 222/249 | 96.5/249 | 0.555 | Complete rebudding ~ duration per 100 min + dataset | 0.62 | 0.33-1.16 | 0.136 | No detectable duration effect on complete rebudding within the sampled *GAL-HO* range. |
| *cdc13-1 yku70Δ* (Ry11047) | 235 | 81/235 | 56.0/235 | 0.871 | Complete rebudding ~ duration per 100 min + dataset | 0.237 | 0.151-0.372 | $4.47 \times 10^{-10}$ | The timing-dependent decline in complete rebudding was retained. |
| *cdc13-1 srs2Δ* (Ry11064) | 198 | 29/198 | 40.3/198 | 0.836 | Complete rebudding ~ duration per 100 min + dataset | 0.248 | 0.140-0.439 | $1.69 \times 10^{-6}$ | The timing-dependent decline was retained in a low-rebudding background. |
| *cdc13-1 sgs1Δ* (Ry10993) | 186 | 71/186 | 49.6/186 | 0.849 | Complete rebudding ~ duration per 100 min + dataset | 0.395 | 0.291-0.537 | $3.19 \times 10^{-9}$ | The timing-dependent decline in complete rebudding was retained. |

1. Complete rebudding was defined as a rebudding score of 2, indicating that both compartments of the arrested mother-daughter pair rebudded. Scores of 0 and 1 were grouped as incomplete rebudding for binary logistic regression.
2. For duration effects, odds ratios represent the change in odds of complete rebudding per additional 100 min of metaphase arrest. Genotype odds ratios compare cdc13-1 mad1Δ with the matched cdc13-1 control. Interaction odds ratios quantify the change in the duration slope associated with MAD1 deletion.
3. Expected counts and AUC values were obtained by applying the original Ry9666 32°C model to the indicated cohort. The AUC reported for the training cohort is an apparent, within-sample estimate.
4. The original Ry9666 34°C cohort comprised datasets cdc13_34_3 to cdc13_34_5 (n = 116). The matched Ry6406 control comprised cdc13_34_1 and cdc13_34_2 (n = 77). These cohorts were not pooled for genotype comparisons.
5. Because only six complete-rebudding events occurred in the original Ry9666 34°C cohort and two component datasets contained no score-2 events, the duration-only refit is reported for that cohort; dataset-adjusted fitting was unstable because of sparse-data separation.
6. The 32°C interaction model included the matched AD104/AD105 experimental block. The 350- and 550-min thresholds were used only for descriptive visualization and were not included in the regression models.

#### Supplementary Table 3. Dataset groups used for model training, model transfer and genotype or lesion-context comparisons

| Dataset group | Strain and experimental context | Cells, *n*[1] | Role in analysis | Main use |
|---|---|---|---|---|
| **Original *cdc13-1*, 32°C** | Ry9666; original reference background | 215 | Primary training cohort | Establish the relationship between metaphase duration and complete rebudding[2] under standard telomere-uncapping conditions |

| | | | | |
|---|---|---|---|---|
| **Original *cdc13-1*, 34°C** | Ry9666; original reference background exposed to more restrictive telomere-uncapping conditions | 116 | Damage-severity model-transfer cohort | Test whether the timing-outcome relationship established at 32°C is retained when telomere dysfunction is increased |
| **Matched *cdc13-1* control, 32°C** | Ry6406; isogenic control for the *mad1Δ* comparison | 100 | Independent experimental cohort | Test transfer of the timing model to a distinct strain and marker background and provide the matched control for the 32°C *MAD1* comparison |
| **Matched *cdc13-1 mad1Δ*, 32°C** | Ry8843; isogenic to Ry6406 except for deletion of *MAD1* | 100 | SAC-genotype comparison | Determine whether loss of *MAD1* changes metaphase duration, baseline rebudding outcome or the duration-outcome relationship at 32°C |
| **Matched *cdc13-1* control, 34°C** | Ry6406; isogenic control for the restrictive-temperature *mad1Δ* comparison | 77 | Severe-damage control cohort | Provide the matched control for evaluating the effect of *MAD1* deletion at 34°C |
| **Matched *cdc13-1 mad1Δ*, 34°C** | Ry8843 | 138 | SAC-genotype comparison under severe damage | Determine whether loss of *MAD1* alters complete rebudding after accounting for metaphase duration under more restrictive conditions |
| ***GAL-HO*** | Ry8847; single inducible HO-generated DNA double-strand break | 249 | Lesion-context comparison and exploratory model transfer | Compare the temporal range and outcome distribution generated by a single defined DNA break with those produced by persistent telomere dysfunction |
| ***cdc13-1 yku70Δ*** | Ry11047 | 235 | DNA-end protection mutant | Test whether the timing-outcome relationship is retained after disruption of Ku-dependent telomere-end protection |
| ***cdc13-1 srs2Δ*** | Ry11064 | 198 | Recombination-control mutant | Test whether altered recombination control changes arrest timing, baseline rebudding competence or the duration-outcome relationship |
| ***cdc13-1 sgs1Δ*** | Ry10993 | 186 | DNA-end processing mutant | Test whether altered helicase and resection functions change arrest timing or the relationship between metaphase duration and rebudding outcome |

**Notes:**

[1] **n** denotes cells with an observed metaphase-exit time and a scorable post-mitotic rebudding outcome that were included in duration-based analyses.
[2] Complete rebudding was defined as a rebudding score of 2, indicating that both compartments of the arrested mother-daughter pair rebudded. Scores of 1 and 0 indicate rebudding of only one compartment and neither compartment, respectively.
[3] The terms “training,” “model transfer” and “comparison” describe the analytical use of each cohort. They do not imply that the independent experimental cohorts constitute external validation studies.

#### Supplementary Videos

**Supplementary Video 1.** Checkpoint adaptation and post-mitotic rebudding in *cdc13-1* cells
Time-lapse imaging of G1-synchronized *cdc13-1* cells (Ry9666) expressing mCherry-Tub1 and Scc1-yEGFP after release at 32°C to induce telomere uncapping. mCherry-Tub1, Scc1-yEGFP and merged channels are shown. Images were acquired every 15 min for 20 h; each displayed frame represents 15 min of experimental time. The movie illustrates checkpoint-induced metaphase arrest, marked by persistent Scc1 (green in merge) and a short bipolar spindle (magenta in merge) followed in adapting cells by Scc1 loss, spindle elongation or disassembly, mitotic exit and post-mitotic rebudding. Scale bar, 5 µm.

**Supplementary Video 2**. Persistent checkpoint arrest in *cdc13-1 cdc5-ad* cells
Time-lapse imaging of G1-synchronized *cdc13-1 cdc5-ad* cells (Ry10642) expressing mCherry-Tub1 and Scc1-yEGFP after release at 32°C to induce telomere uncapping. mCherry-Tub1, Scc1-yEGFP and merged channels are shown. Two representative cells are shown. Images were acquired every 15 min for 20 h; each displayed frame represents 15 min of experimental time. The movie illustrates sustained checkpoint arrest, characterized by persistent Scc1 (green in merge) and short bipolar spindles (magenta in merge). Scale bar, 5 µm.

**Supplementary Video 3.** Mitotic progression in checkpoint-deficient *cdc13-1 rad9Δ* cells
Time-lapse imaging of G1-synchronized *cdc13-1 rad9Δ* cells (Ry10990) expressing mCherry-Tub1 (magenta) and Scc1-yEGFP (green) after release at 32°C to induce telomere uncapping. Merged channels are shown. Images were acquired every 15 min for 20 h; each displayed frame represents 15 min of experimental time. Two representative cells are shown. The movie illustrates continued mitotic progression and successive budding events in the absence of a sustained *RAD9*-dependent checkpoint arrest, despite persistent telomere dysfunction. Scale bar, 5 µm.

**Supplementary Video 4.** Delayed and asynchronous checkpoint establishment in *cdc13-1 exo1Δ* cells
Time-lapse imaging of G1-synchronized *cdc13-1 exo1Δ* cells (Ry10998) expressing mCherry-Tub1 (magenta) and Scc1-yEGFP (green) after release at 32°C to induce telomere uncapping. Merged channels are shown. Images were acquired every 15 min for 20 h; each displayed frame represents 15 min of experimental time. The movie illustrates delayed checkpoint engagement: cells initially progress through mitosis, whereas later lineage events show increasingly prolonged metaphase. Scale bar, 5 µm.

#### **Supplementary Method:** Predictive modelling, validation, cross-context transfer and refitting of the metaphase-duration outcome model

##### Overview of the modelling strategy

The modelling analysis tested whether metaphase arrest duration predicted post-adaptation proliferative outcome across independent datasets, damage severities and genetic backgrounds within the *cdc13-1* telomere-dysfunction system. Application of the model to *GAL-HO* cells were analyzed separately as an exploratory cross-context transfer analysis. This analysis assessed whether a predictor derived from persistent telomere dysfunction retained calibration or outcome-ranking performance in a distinct lesion context; *GAL-HO* was not treated as an independent validation cohort. Metaphase duration was defined as the interval between bipolar spindle formation and cohesin cleavage, marked by loss of Scc1 signal or by bipolar spindle formation and disassembly. Post-adaptation proliferative outcome was scored according to rebudding of the two compartments of the arrested mother-daughter pair: score 2, both compartments rebudded; score 1, one compartment rebudded; score 0, neither compartment rebudded.
For the primary logistic-regression analysis, cells with score 2 were classified as showing full proliferative re-entry, whereas cells with score 0 or 1 were classified as incomplete or failed proliferative re-entry. Metaphase duration was scaled to 100-min intervals so that odds ratios represent the change in odds of full proliferative re-entry for each additional 100 min spent in metaphase arrest.

The primary *cdc13-1* training model was fit to the original *cdc13-1* datasets, with dataset included as a batch term to account for experiment-to-experiment variability. For validation within telomere-dysfunction backgrounds, the original *cdc13-1* model was applied to independent datasets to estimate the predicted probability of full proliferative re-entry in each cell. Because dataset-specific batch terms are internal to the training datasets, external prediction was evaluated by model discrimination and calibration, and each external group was then refit directly to test whether metaphase duration remained predictive in that condition. The same prediction metrics were calculated for *GAL-HO* cells solely to quantify cross-context model transfer. Because *GAL-HO* cells sampled a restricted arrest-duration range and showed a high frequency of full proliferative re-entry, these metrics were interpreted as defining the limits of transfer rather than as validation of the original model.

Model performance after external prediction was summarized using the observed number of fully recovered cells, the expected number of fully recovered cells predicted by the original *cdc13-1* model, and the area under the receiver operating characteristic curve. Where available, Brier score and classification accuracy at a 0.5 probability threshold were also retained. Calibration and discrimination were interpreted separately: differences between observed and expected proliferative re-entry indicated imperfect calibration, whereas AUC assessed whether the model retained outcome-ranking performance.

For genotype comparisons, combined logistic-regression models included metaphase duration, genotype and dataset or block where appropriate. Genotype terms tested whether proliferative re-entry differed between genotypes after accounting for metaphase duration. Metaphase duration x genotype interactions

tested whether the slope of the timing-outcome relationship differed between genotypes.
The 350-min and 550-min timing thresholds were used only for visualization and descriptive stratification. They were not used for primary statistical inference.

**Primary model:** logit[P(full proliferative re-entry)] = $\beta_0$ + $\beta_1$(metaphase duration/100 min) + $\beta_2$(dataset).
**Genotype-comparison model:** logit[P(full proliferative re-entry)] = $\beta_0$ + $\beta_1$(metaphase duration/100 min) + $\beta_2$(genotype) + $\beta_3$(dataset/block) + $\beta_4$(metaphase duration x genotype).
